# Off-Target Structural Insights: ArnA and AcrB in Bacterial Membrane Protein Cryo-EM Analysis

**DOI:** 10.1101/2025.05.31.656883

**Authors:** Mehmet Caliseki, Ufuk Borucu, Sathish K. N. Yadav, Christiane Schaffitzel, Burak Veli Kabasakal

## Abstract

Membrane protein quality control in *Escherichia coli* involves coordinated actions of the AAA+ protease FtsH, the insertase YidC, and the regulatory complex HflKC. These systems maintain proteostasis by facilitating membrane protein insertion, folding, and degradation. To gain structural insights into a putative complex formed by FtsH and YidC, we performed single-particle cryogenic electron microscopy on detergent-solubilized membrane samples, from which FtsH and YidC were purified using Ni-NTA affinity and size exclusion chromatography. Although SDS-PAGE analysis indicated a high purity of these proteins, cryo-EM datasets unexpectedly yielded high-resolution structures of ArnA and AcrB at 4.0 Å and 2.9 Å resolutions, respectivelyArnA is a bifunctional enzyme involved in lipid A modification and polymyxin resistance, while AcrB is a multidrug efflux transporter of the AcrAB–TolC system. ArnA and AcrB are known membrane protein contaminants after Ni-NTA purifications. However, they were also detected in Strep-Tactin affinity purified FtsH-YidC samples. ArnA, typically cytoplasmic, was consistently found in membrane-isolated samples, indicating an association with membrane components. Two-dimensional class averages revealed additional particles resembling GroEL, cytochrome bo₃ oxidase, and the AAA+ cytoplasmic domain of FtsH. Although our preparation was initially considered homogeneous, neither full-size FtsH nor the intact FtsH–YidC complex could be resolved. This case highlights that cryo-EM can reveal structural information beyond the intended target, and that overexpression of membrane proteins involved in quality control may lead to the co-purification of proteins such as ArnA and AcrB.

## 1. INTRODUCTION

Bacterial membrane proteostasis relies on the coordinated action of membrane protein biogenesis and degradation pathways, which ensure the proper insertion, folding, and removal of membrane proteins (Akiyama, 2009; Watkins et al., 2022; Njenga et al., 2023). Membrane protein quality control plays a central role in this system by preventing the accumulation of misfolded or non-functional proteins that could compromise membrane integrity and disrupt cellular function (Akiyama, 2009; Dalbey et al., 2012; G. Liu et al., 2020; Njenga et al., 2023; Watkins et al., 2022; Ero et al., 2024).

In gram-negative bacteria, inner membrane protein quality control involves distinct systems responsible for protein insertion and degradation. The membrane-bound AAA+ protease FtsH removes misfolded or unassembled membrane proteins in an ATP- and Zn²⁺-dependent manner (Akiyama, 2002; Ito & Akiyama, 2005; Bieniossek et al., 2006; Langklotz et al., 2012; Carvalho et al., 2021). FtsH forms a complex with the membrane proteins HflKC, which modulate its proteolytic activity and substrate specificity (Kihara et al., 1998; Saikawa et al., 2004; Qiao et al., 2022; Ma et al., 2022; Akkulak et al., 2024; Ghanbarpour et al., 2025). In parallel, the insertase YidC facilitates the membrane integration and proper folding of nascent polypeptides, functioning either independently or in association with the Sec translocon (Houben et al., 2000; Laan et al., 2005; Beck et al., 2001; Schulze et al., 2014; Dalbey et al., 2014; Komar et al., 2016; Botte et al., 2016; Petriman et al., 2018; Shanmugam et al., 2019; Caliseki et al., 2025). These systems act together to maintain membrane proteostasis and prevent the accumulation of defective protein species under both normal and stress conditions (Van Bloois et al., 2008; Akiyama, 2009; Njenga et al., 2023; Caliseki et al., 2025).

To gain structural insight into FtsH–YidC interactions, single particle cryogenic electron microscopy (cryo-EM) was performed on detergent-solubilized, affinity-purified membrane proteins. FtsH-YidC samples were purified using Ni-NTA affinity chromatography followed by size exclusion chromatography (SEC). SDS-PAGE analysis showed a very high enrichment of FtsH and YidC in the purified fractions, indicating that the two proteins interact and form a complex (Sup. Fig 1-3).

Although the primary objective was to determine the structure of the FtsH–YidC complex, single-particle cryo-EM analysis unexpectedly yielded high-resolution reconstructions of two unrelated proteins: ArnA and AcrB. ArnA is a bifunctional enzyme that catalyzes the biosynthesis of 4-amino-4-deoxy-L-arabinose (L-Ara4N), a nucleotide-activated sugar that is incorporated into lipid A to reduce its net negative charge and thereby enhance resistance to cationic antimicrobial peptides such as polymyxins (Gatzeva-Topalova et al., 2005; Williams et al., 2005; Yang et al., 2019). AcrB is a key component of the AcrAB–TolC multidrug efflux system, responsible for exporting diverse substrates across the bacterial envelope (Murakami et al., 2002; Seeger et al., 2006; Takatsuka et al., 2010; Trinh et al., 2023). ArnA and AcrB are known contaminants in Ni-NTA affinity purification (Bolanos-Garcia & Davies, 2006; Veesler et al., 2008; Glover et al., 2011; Andersen et al., 2013). However, we also detected these proteins in Strep-Tactin purified fractions by mass spectrometry, suggesting that their presence may reflect a specific association rather than nonspecific contamination (Caliseki et al., 2025)

Although ArnA is typically classified as a cytoplasmic protein, it was consistently detected in membrane-isolated fractions, suggesting a potential association with membrane-localized assemblies (Caliseki et al., 2025). In addition to ArnA and AcrB, two-dimensional (2D) class averages revealed distinct particles resembling GroEL (Fujita et al., 2023), cytochrome bo₃ oxidase (Su et al., 2021), and the AAA+ cytoplasmic domain of FtsH (Lee et al., 2011; Qiao et al., 2022); the presence of these proteins was confirmed by mass spectrometry (Sup. Table 1). Although the sample was initially considered homogeneous based on purification and SDS PAGE analysis, the presence of multiple co-purified proteins in the cryo-EM data revealed unexpected compositional complexity. Neither full-size FtsH nor the FtsH–YidC complex could be resolved, also YidC alone could not be detected in the cryo-EM data set. Two unrelated protein complexes were detected with considerable abundance and purity in the cryo-EM preparations, allowing to solve high-resolution structures. This unexpected outcome shows that cryo-EM can uncover hidden structural information in membrane protein preparations (Su et al., 2021), providing insight into biologically relevant proteins beyond the intended target, highlighting the pitfalls as well as sensitivity and capability of this technique.

## 2. MATERIAL & METHODS

### 2.1. Expression Systems and Plasmid Design

The FYHC Co-T expression system was used to co-express FtsH, YidC, HflK, and HflC in *Escherichia coli* C43 (DE3) cells (Caliseki et al., 2025). This system employed the pETDuet-1 (Novagen, 71146-3) and pRSFDuet-1 (Novagen, 71341) plasmids, both driven by T7 promoters. The pETDuet-1 vector encoded YidC with a C-terminal 10×His tag and FtsH with a C-terminal 3×StrepTag II, while the pRSFDuet-1 plasmid carried untagged HflK and HflC. Ampicillin and kanamycin resistance markers enabled co-transformation and stable maintenance of both plasmids for simultaneous expression of the target proteins.

### 2.2. Membrane Protein Expression, Isolation and Purification

*Escherichia coli* C43 (DE3) cells harbouring the FYHC Co-T expression system were used for recombinant protein expression. Starter cultures were grown overnight at 37□°C in Luria-Bertani (LB) medium containing appropriate antibiotics. A 5□mL aliquot was transferred into 2□L baffled flasks containing 500□mL of Terrific Broth (TB; Sigma-Aldrich, T0918). Typical culture volumes were 6L. Cultures were incubated at 37□°C until an optical density at 600□nm (OD₆₀₀) of ∼0.8 was reached. Protein expression was induced by addition of 0.4□mM isopropyl-β-D-thiogalactopyranoside (IPTG), and cultures were incubated at 18□°C for 18□hours.

Cells were harvested by centrifugation at 5,000□×□g for 30□min at 4□°C and resuspended in lysis buffer (20□mM HEPES, pH□7.5; 150□mM NaCl; 8% glycerol; 2□mM MgCl₂) supplemented with 1□mM phenylmethylsulphonyl fluoride (PMSF; Roche, 11359061001), one tablet of EDTA-free protease inhibitor cocktail (Pierce, A32965), and 1□µg/mL benzonase nuclease (Millipore, E1014). The cell pellet was resuspended and incubated for 1 hour at 4□°C with gentle agitation to allow complete resuspension and enzyme activity. Cells were then lysed by sonication on ice using a probe sonicator at 40% amplitude, with 2□seconds on / 8□seconds off cycles for 2□minutes, repeated twice. The lysate was clarified by centrifugation at 10,000□×□g for 1.5□hours at 4□°C.

Membrane fractions were isolated from the lysate by ultracentrifugation at 100,000□×□g for 1□hour at 4□°C. The resulting pellets were resuspended in a solubilization buffer composed of 20□mM HEPES (pH□7.5), 150□mM NaCl, 8% (v/v) glycerol, and 2□mM MgCl₂, supplemented with detergents. For the first and second cryo-EM screenings, 1% (w/v) n-dodecyl-β-D-maltoside (DDM, Anatrace, Cat# D310A) was added to the solubilization buffer. For the third screening, the buffer was further supplemented with 0.1% (w/v) cholesterol hemisuccinate (CHS, Anatrace, Cat# CH210) in addition to 1% DDM to reduce aggregation and enhance membrane protein stability. Solubilization was performed at 4□°C for 1□hour with gentle agitation.

Subsequently, solubilized membrane proteins were chemically crosslinked by adding 0.25□mM DSP for the first Cryo-EM screening, or 1□mM DSP for the second and third screenings, followed by incubation at 4□°C for 2□hours. The reaction was quenched by addition of 20□mM Tris-HCl (pH□7.5) and incubated on ice for 15□minutes. Insoluble material was removed by ultracentrifugation at 100,000□×□g for 1□hour at 4□°C. The resulting supernatant was clarified by filtration through a 0.45□µm membrane prior to affinity purification.

Solubilized and crosslinked membrane proteins were first subjected to Ni-NTA affinity purification using the gravity flow technique. For the first and second cryo-EM screenings, purification buffers contained 0.02% (w/v) DDM. For the third screening, buffer composition was modified to include 0.05% (w/v) DDM and 0.005% (w/v) CHS to improve protein stability. Size-exclusion chromatography (SEC) was subsequently performed on an ÄKTA micro FPLC system (Cytiva) using a Superose 6 Increase 10/300 GL column equilibrated in the corresponding detergent-containing buffer.

Ni-NTA resin (HisPur Superflow Agarose, Thermo Scientific, 25215) was equilibrated with binding buffer composed of 20□mM HEPES (pH□7.5), 150□mM NaCl, 8% (v/v) glycerol, 2□mM MgCl₂, the appropriate concentration of DDM, and 10□mM imidazole. After protein binding, the resin was washed sequentially with high-salt buffer (including 1□M NaCl) and wash buffer containing 50□mM imidazole. Bound proteins were eluted using elution buffer containing 300□mM imidazole.

Eluted fractions were collected and diluted 3:1 with imidazole-free buffer (20□mM HEPES, pH□7.5; 150□mM NaCl; 8% glycerol; 2□mM MgCl₂; appropriate DDM concentration) to reduce the imidazole concentration prior to concentration. The diluted samples were then concentrated using Amicon® Ultra-15 Centrifugal Filter Units (100□kDa MWCO; Merck Millipore, UFC910024) at 4□°C by centrifugation at 4,000□×□g using a swinging-bucket rotor. Concentration was continued until the final volume reached approximately 1.5□mL with a protein concentration of ∼6-8□mg/mL.

Concentrated samples were subjected to size-exclusion chromatography (SEC) using an ÄKTA™ micro system (Cytiva) equipped with a Superose 6 Increase 3.2/300 column (Cytiva, Cat# 29091598), pre-equilibrated with buffer containing 20□mM HEPES (pH□7.5), 100□mM NaCl, and 2□mM MgCl₂. For the first and second cryo-EM screenings, the SEC buffer contained 0.02% (w/v) DDM. For the third sample, the buffer was supplemented with 0.05% (w/v) DDM and 0.005% (w/v) CHS. A total of 50□µL sample was injected per run, and separation was carried out at a flow rate of 0.05□mL/min.

### 2.3. SDS-PAGE and Western Blot Analysis

Protein expression and purification profiles were assessed by SDS–PAGE (Sup. Fig. 1-3) and Western blotting (Sup. Fig. 1). Electrophoresis was performed using commercial 4–12% Bolt™ Bis-Tris Plus Mini Gels (Thermo Fisher Scientific, NW04125BOX) at 180□V in Tris– glycine running buffer (25□mM Tris, 192□mM glycine, 0.1% SDS, pH□8.3). After separation, proteins were visualized by Coomassie Brilliant Blue staining.

For Western blotting, proteins were transferred from the SDS gels onto 0.2□µm nitrocellulose membranes (Bio-Rad) using the Trans-Blot Turbo system (Bio-Rad) at 25□V for 7□minutes. Membranes were blocked with 3% (w/v) BSA in TBS-T (20□mM Tris-HCl pH□7.5, 150□mM NaCl, 0.1% v/v Tween-20) for 1□hour at room temperature. Tagged proteins were detected using HRP-conjugated primary antibodies: anti-His (Qiagen, 1:5000), anti-StrepTag II (Sigma-Aldrich, 1:5000), and anti-Flag (Rockland, 1:10000). Following antibody incubation and washing, chemiluminescence signals were developed with Pierce™ ECL substrate (Thermo Scientific) (Sup. Fig. 1).

### 2.4. Negative Stain Electron Microscopy

SEC-purified samples were diluted into SEC buffer to final protein concentrations of approximately 0.05□mg/mL, 0.025□mg/mL, or 0.01□mg/mL. A 4□µL aliquot was applied onto glow-discharged carbon-coated copper grids (CF300-CU, 300 mesh; Electron Microscopy Sciences) and incubated for 1□minute at room temperature. Excess sample was blotted off with filter paper.

Grids were stained with 2% (w/v) uranyl acetate for 1□minute, followed by blotting with filter paper. A second drop of uranyl acetate was applied to wash the grid, and excess stain was again removed with filter paper. Grids were left to air-dry completely before imaging.

NS-EM images were acquired using a Tecnai T12 transmission electron microscope (Thermo Fisher Scientific) operated at 120□kV equipped with a Ceta 16 M CCD detector at the Wolfson Bioimaging Facility (University of Bristol), at magnifications ranging from 25,000× to 86,000×.

### 2.5. Cryo-EM Grid Preparation, Screening and Data Collection

Cryo-EM grids were prepared using SEC-purified samples obtained under three different preparation conditions (see above). In the first screening, grids were vitrified using a Leica EM GP2 system, while in the second and third screenings, grids were vitrified using a Vitrobot Mark IV system (Thermo Fisher Scientific). Prior to sample application, grids were rendered hydrophilic using an Elmo glow discharge unit (Cordouan Technologies) at a setting of 300□mC. A 4□µL aliquot of each sample was applied to glow-discharged R2.2/2□nm carbon-coated 300-mesh copper grids (Quantifoil). The initial cryo-EM grid preparation involved grids vitrified using the Leica EM GP2 system and screened on a Talos Arctica, with samples crosslinked using 0.25 mM DSP crosslinker. The second and third cryo-EM screenings employed grids with 1 mM DSP crosslinked samples vitrified using the Vitrobot Mark IV system under the following conditions: 4□°C, 100% relative humidity, blot force of 0 or 1, 10□s wait time, 2□s blot time, no drain time, and back-side blotting enabled. For the first dataset from the second screening, 0.4% w/v CHAPS was added prior to vitrification to reduce aggregation. In the second dataset from the third screening, 0.05% w/v DDM was used with 0.005% w/v CHS maintained in the final SEC buffer.

Cryo-EM grid screening and data collection were conducted in the GW4 Facility for High-Resolution Electron Cryo-Microscopy at the University of Bristol using a Talos Arctica transmission electron microscope (Thermo Fisher Scientific) operating at 200□kV equipped with an energy filter and a K2 direct electron detector (Gatan). No data was collected from the first screening due to severe sample aggregation (not shown). The first dataset (3,908 movies) and second dataset (7,506 movies) were obtained from the second and third grid screenings, respectively. Both datasets were recorded at 130,000× nominal magnification with a calibrated pixel size of 1.05□Å, 6-second exposure per movie (60 frames), and total electron doses of 58.2□e⁻/Å² and 60.6□e⁻/Å², respectively (Table 1).

**Table 1.**
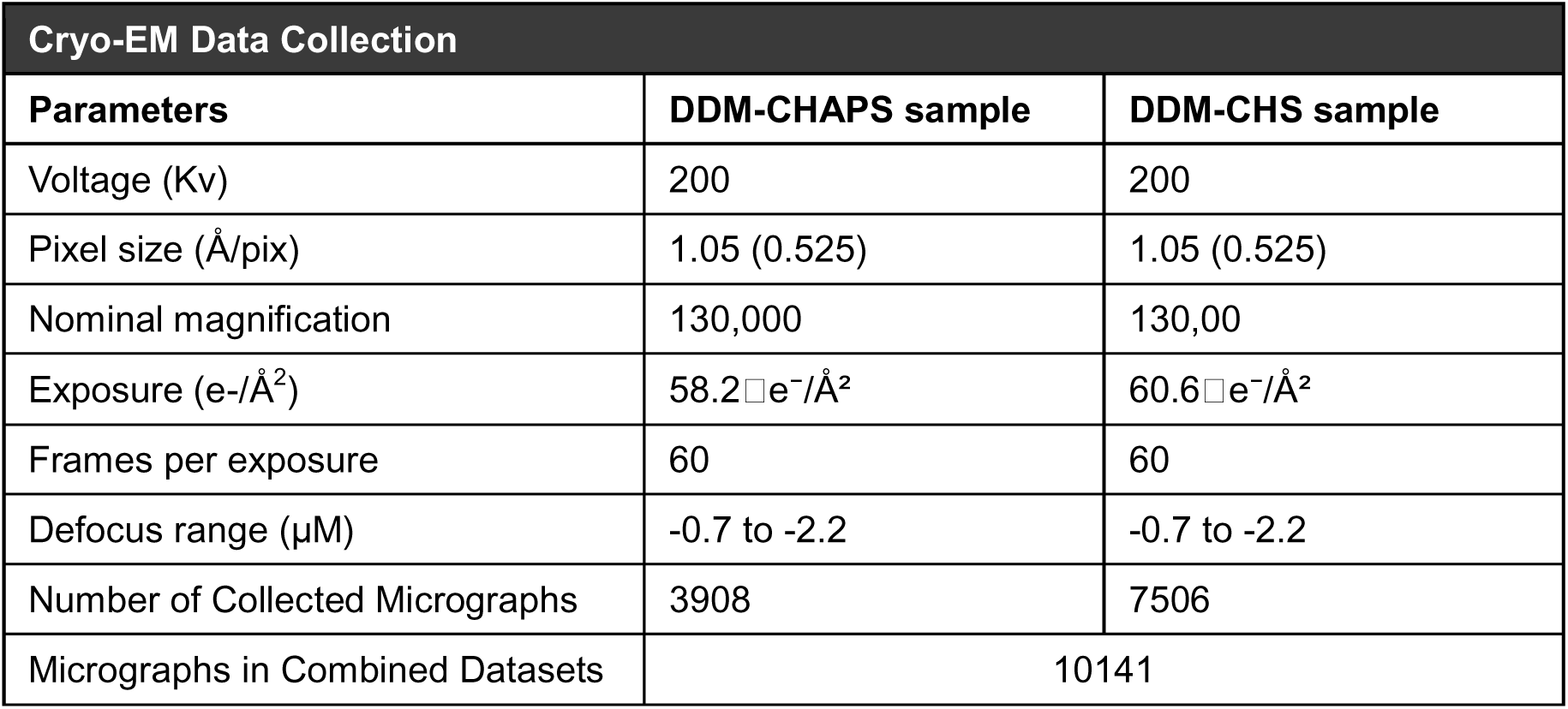
Cryo-EM Data Collection and Initial Data Processing.

### 2.5. Cryo-EM Data Processing

Cryo-EM image processing was performed using cryoSPARC (Punjani et al., 2017). Movies from both datasets were initially processed independently (Figure 1). Patch motion correction and patch CTF estimation (Downing & Glaeser, 2008; Punjani et al., 2017) were performed separately, followed by micrograph curation.

**Figure 1.**
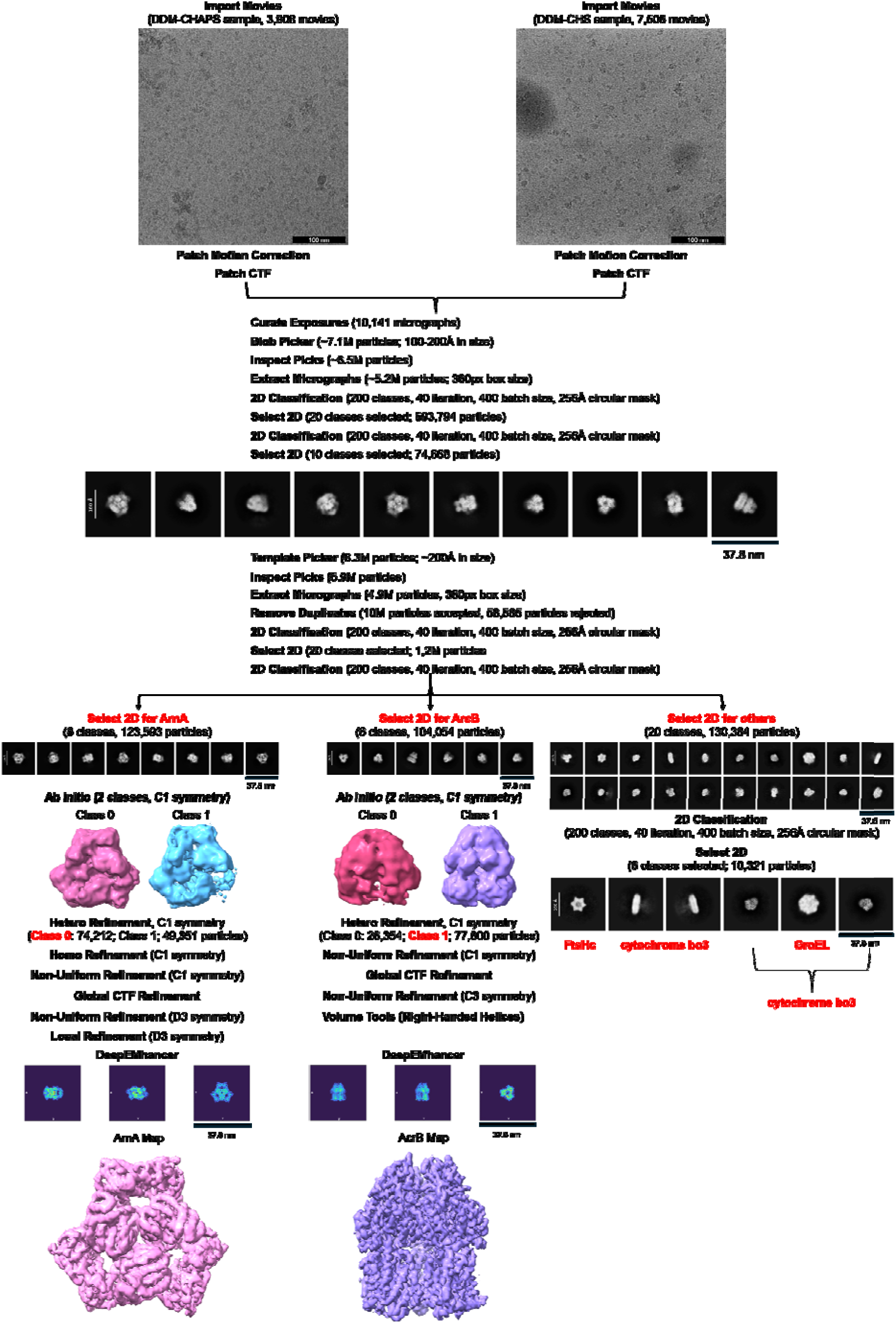
Optimized cryoSPARC workflow using the combined Cryo-EM dataset. Two cryo-EM datasets were collected from DSP-crosslinked membrane protein samples. The first dataset (3,908 movies) was prepared with 0.02% DDM and 0.4%w/v CHAPS as an additive, and the second dataset (7,506 movies) with 0.05% DDM and 0.005% w/v cholesterol hemisuccinate (CHS). Motion correction and CTF estimation were performed separately for each dataset. 10,141 micrographs were used for particle picking resulting in 7.13 million particles (estimated diameter: 100–200CÅ). Two rounds of 2D classification (200 classes, 256CÅ mask, 40 online-EM iterations, batch size 400) retained 74,668 particles [scale bar: 37.8 nm]. These classes were used as templates for template-based picking, which yielded 6.3 million particles. After re-extraction and duplicate removal, a final set of 10.13 million unique particles was obtained. 2D classification yielded 1.26 million particles distributed across the best 20 classes. For ArnA, 123,593 particles corresponding to ArnA 2D classes [scale bar: 37.8 nm for 2D class averages] were selected and processed through *ab-initio* reconstruction with C1 symmetry, followed by heterogeneous and non-uniform refinement using D3 symmetry. For AcrB, 104,054 particles corresponding to AcrB 2D classes [scale bar: 37.8 nm for 2D class averages] were selected and refined using C3 symmetry. Global CTF refinement and DeepEMhancer post-processing were applied to both maps. Handedness correction for AcrB was performed using the Volume Tools utility in cryoSPARC. Particles corresponding to GroEL, cytochrome bo₃ oxidase, and other minor components were also detected during classification [scale bar: 37.8 nm for 2D class averages].

For the first dataset, particle picking in cryoSPARC (Punjani et al., 2017) yielded 671,626 particles, of which 44,881 were retained after 2D classification (Sup. Fig. 4A). For the second dataset, 2,123,935 particles were extracted using the same 360-pixel box size. After 2D classification, 185,213 particles were retained (Sup. Fig. 4B).

As similar 2D class averages were observed in both datasets, the micrographs were merged at the CTF refinement stage for joint downstream processing. From the combined dataset, 10,141 curated micrographs were selected. Initial particle picking was performed using blob-based algorithms with estimated particle diameters between 100 and 200□Å, yielding approximately 7.13 million particles. After particle picking, low-quality particles were removed based on 2D class averages, yielding a curated dataset of 6.57 million particles.

Subsequently, two rounds of 2D classification were then performed using a circular mask diameter of 256 Å. The first round yielded 593,794 particles across 20 selected classes. The second round refined this to 74,668 particles from 10 well-defined classes, which were then used for template-based particle picking. This step identified 6.3 million particles. After combining with the initial blob-picked set and removing duplicates, 10.13 million unique particles were retained. A final round of 2D classification resulted in a cleaned dataset containing 1.26 million particles for downstream processing (Figure 1).

2D class averages revealed the presence of multiple co-purified protein species, including ArnA, AcrB, GroEL, cytochrome bo₃ oxidase, and FtsH. Among the total dataset, 130,384 particles were assigned to classes not matching ArnA or AcrB. These particles were further analyzed by 2D classification to group them based on structural similarity, using the same parameters described previously.

For ArnA, a total of 123,593 particles were selected and processed using *ab initio* reconstruction followed by heterogeneous, homogeneous and non-uniform refinement in cryoSPARC with C1 symmetry. In the final stage, D3 symmetry was imposed during the second non-uniform refinement (Punjani et al., 2020). The resulting map was sharpened using DeepEMhancer (Sanchez-Garcia et al., 2021).

For AcrB, 104,054 particles were subjected to the same processing workflow. Initial *ab initio* reconstruction and subsequent heterogeneous and non-uniform refinement were performed with C1 symmetry. C3 symmetry was applied during the final non-uniform refinement step (Punjani et al., 2020). Handedness correction was conducted using volume tools within cryoSPARC. The final map was sharpened using DeepEMhancer (Sanchez-Garcia et al., 2021).

The Fourier Shell Correlation (FSC) curves for ArnA and AcrB were calculated using Phenix Mtriage (Afonine et al., 2018; Liebschner et al., 2019), based on the final map and the half-maps generated in cryoSPARC.

### 2.6. Model Building and Validation

Atomic model building and validation of the ArnA and AcrB cryo-EM maps were performed using an integrated modeling workflow. The initial fitting of atomic models into the experimental maps was carried out using MolRep (Vagin & Teplyakov, 2010) within the CCP-EM suite (Agirre et al., 2023). Template structures PDB: 6PIH for ArnA (Yang et al., 2019) and PDB: 7RR7 for AcrB (Trinh et al., 2023) were used for molecular replacement.

Subsequent real-space refinement was performed using Phenix (Liebschner et al., 2019), incorporating local grid sampling, map-weighted geometry restraints, and secondary structure preservation. To improve global stereochemistry and atomic displacement parameters, an additional round of refinement was conducted using Refmac as implemented in CCP-EM (Burnley et al., 2017). Manual model adjustment, including side-chain placement, rotamer correction, and loop rebuilding, was carried out in Coot (Emsley et al., 2010).

All final models were validated using Phenix’s comprehensive validation tools, including Molprobity scores, Ramachandran outlier detection, and map-to-model correlation metrics (Liebschner et al., 2019) (Table 2).

**Table 2.**
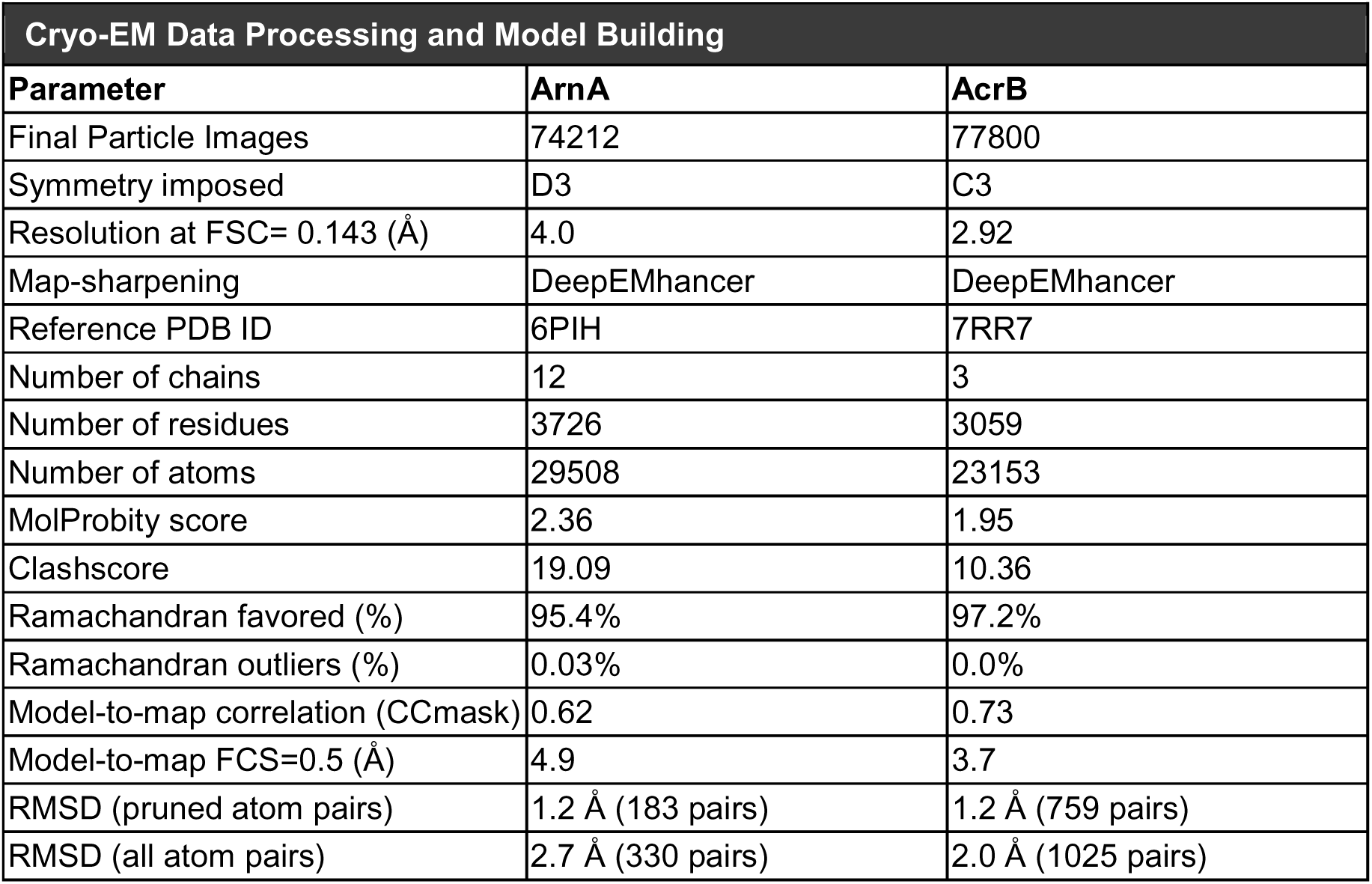
Cryo-EM Data Processing and Protein Modelling for ArnA and AcrB.

| Cryo-EM Data Processing and Model Building |  |  |
| --- | --- | --- |
| Parameter | ArnA | AcrB |
| Final Particle Images | 74212 | 77800 |
| Symmetry imposed | D3 | C3 |
| Resolution at FSC= 0.143 (Å) | 4.0 | 2.92 |
| Map-sharpening | DeepEMhancer | DeepEMhancer |
| Reference PDB ID | 6PIH | 7RR7 |
| Number of chains | 12 | 3 |
| Number of residues | 3726 | 3059 |
| Number of atoms | 29508 | 23153 |
| MolProbity score | 2.36 | 1.95 |
| Clashscore | 19.09 | 10.36 |
| Ramachandran favored (%) | 95.4% | 97.2% |
| Ramachandran outliers (%) | 0.03% | 0.0% |
| Model-to-map correlation (CCmask) | 0.62 | 0.73 |
| Model-to-map FCS=0.5 (Å) | 4.9 | 3.7 |
| RMSD (pruned atom pairs) | 1.2 Å (183 pairs) | 1.2 Å (759 pairs) |
| RMSD (all atom pairs) | 2.7 Å (330 pairs) | 2.0 Å (1025 pairs) |

## 3. RESULTS

### 3.1. Initial Cryo-EM Screening Reveals Aggregation in FtsH-YidC Samples without Additives

The first cryo-EM screening was carried out to investigate whether the sample was suitable for structural analysis of a potential FtsH-YidC complex. Membranes were solubilized in 1% n-dodecyl-β-D-maltoside (DDM) and crosslinked with 0.25□mM dithiobis(succinimidyl propionate) (DSP), followed by Ni-NTA affinity purification and size-exclusion chromatography (SEC) without the inclusion of stabilizing additives. Western blot analysis confirmed the presence of His-tagged YidC and Strep-tagged FtsH in SEC fractions (Sup. Fig. 1A-C). The major SEC peak eluting at approximately 1.6□mL was collected for further analysis, and SDS-PAGE confirmed enrichment of the target proteins in this fraction (Sup. Fig.□1D-E). Negative-stain electron microscopy revealed a homogeneous distribution of particles without obvious aggregation (Sup. Fig. 1F). In contrast, cryo-EM grids vitrified using a Leica EM GP2 system and screened on a Talos Arctica microscope exhibited extensive particle aggregation within the vitreous ice, preventing further structural analysis (not shown). The observed clustering in cryo-EM micrographs could be attributed to vitrification conditions, prompting further optimization of this step in subsequent preparations.

### 3.2. CHAPS Improves Particle Dispersion in Crosslinked FtsH-YidC Samples

A second cryo-EM screening was performed after SEC purification of the DSP-crosslinked FtsH–YidC sample (Sup. Fig. □2 and 4A). In this preparation, 0.4% (w/v) CHAPS was included to enhance particle dispersion and reduce aggregation. As a zwitterionic detergent, CHAPS was used to stabilize membrane protein assemblies by minimizing hydrophobic interactions that typically lead to aggregation during grid preparation (Kampjut et al., 2021). Compared to the first screening, this approach resulted in improved vitrification quality, as evidenced by reduced aggregation and more homogeneous particle distribution (Figure 1 and Sup. Fig. □4A). A total of 3,908 micrographs were collected for this dataset. Despite improved sample quality, the number of usable particles and their angular distribution remained insufficient for high-resolution reconstruction (Sup. Fig. 4B), indicating the need for further cryo-EM data collection.

### 3.3. Enhanced Cryo-EM Grid Quality Achieved through DDM–CHS Solubilization

A final cryo-EM screening was performed using the C6 fraction obtained after SEC purification of the DSP-crosslinked FtsH–YidC sample (Sup. Fig.□3B and 4C). In this experiment, membrane proteins were solubilized using a buffer containing 1% w/v DDM and 0.1% w/v CHS. CHS was included to mimic membrane cholesterol and enhance the structural stability of membrane proteins during detergent solubilization and vitrification (Li, 2022). Following solubilization, all samples and buffers were supplemented with 0.05% (w/v) DDM and 0.005% (w/v) CHS. This preparation yielded grids with significantly reduced aggregation and well-dispersed particles (Sup. Fig. 4C). A total of 7,506 micrographs were collected for this dataset and processed in cryoSPARC. Following multiple rounds of 2D classification, 185,213 particles were selected (Sup. Fig. 4D).

Based on the high similarity observed in the 2D class averages, micrographs from this preparation and the previous CHAPS-containing sample were combined after Patch CTF estimation. The combined dataset was then used to select high-quality trimeric and tetrameric particle classes for subsequent 3D reconstruction.

### 3.4. Off-target Structural Determination of ArnA and AcrB in FtsH-YidC Cryo-EM Dataset

Cryo-EM analysis was performed on two independently collected datasets derived from crosslinked FtsH-YidC complex samples. Although the samples were optimized to preserve native interactions, the expected FtsH–YidC structure could not be resolved. Instead, dominant particle populations corresponding to ArnA and AcrB were consistently identified during data processing.

An optimized cryoSPARC processing pipeline was established and followed for all subsequent analyses (Figure□1). In the 2D class averages, distinct particle classes resembling the cytoplasmic domains of FtsH, GroEL, and cytochrome bo₃ oxidase were also observed alongside ArnA and AcrB, but they were not processed further for 3D reconstruction due to low particle numbers and preferred views.

To achieve high-resolution reconstructions, *ab-initio* reconstruction and refinement were performed for ArnA and AcrB. For ArnA, 123,593 particles were processed through heterogeneous and non-uniform refinement, initially using C1 symmetry and later D3 symmetry. The final reconstruction yielded a map with a global resolution of 4.0 Å, as determined by the FSC_0.143_ criterion (Figure□2 and Sup. Fig. 5).

For AcrB, 104,054 particles were refined using C1 symmetry, followed by C3 symmetry in the final non-uniform refinement step, resulting in a cryo-EM map with a global resolution of 2.92□Å, as determined by the FSC_0.143_ criterion (Figure 3 and Sup. Fig 6). Global CTF refinement and DeepEMhancer post-processing were applied. Handedness correction was performed using cryoSPARC volume tools. Local resolution estimation and directional FSC analysis were performed using Phenix, confirming moderate anisotropy and high overall map quality for both ArnA and AcrB reconstructions (Figure 1).

### 3.5. High-Resolution Cryo-EM Structure and Model Validation of Hexameric ArnA

A structural model of ArnA was built using PDB: 6PIH as a starting model by molecular replacement using MolRep (Vagin & Teplyakov, 2010) within the CCP-EM interface (Burnley et al., 2017) (Figure 2A). Refinement was performed using Phenix Real Space Refinement, followed by optimization in Refmac and final manuel adjustments in Coot to complete the atomic model (Figure 2B). The final model was validated using Phenix tools, which included both geometry and density correlation analyses (Sup. Fig 5A).

**Figure 2.**
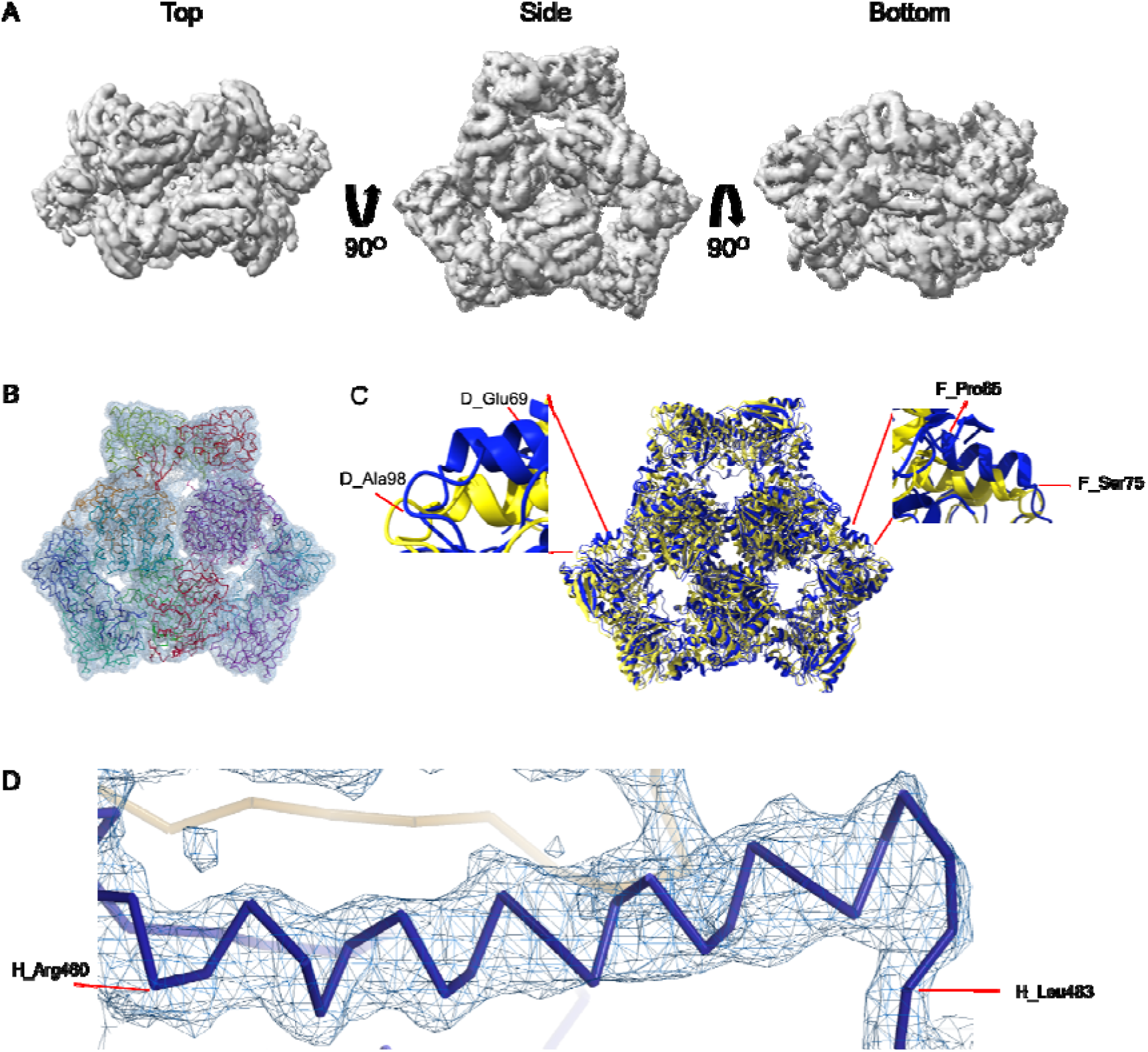
Cryo-EM density map and model fitting of the hexameric ArnA complex. (A) Cryo-EM density map of the ArnA hexamer reconstructed at 4.0LJÅ resolution, shown from three orientations: top, side, and bottom views. The map reveals the characteristic two-layered architecture of ArnA and clearly resolved secondary structure elements. (B) Final atomic model of ArnA fitted into the cryo-EM density map. Each of the 12 subunits is displayed in a different color to illustrate the hexameric arrangement. (C) Structural alignment of the final cryo-EM model (blue) with the reference crystal structure (yellow) using ChimeraX. The alignment yielded an RMSD of 1.2LJÅ. Zoomed-in views show local conformational deviations in loop regions: Glu69-Ala98 in Chain D (left) and Pro65-Ser75 in Chain F (right). (D) Close-up view showing the model-to-map fit for a peptide segment (Leu483–Arg460). The density mesh is contoured at 1.7σ, showing clear peptide backbone density and supporting accurate model placement.

Local resolution estimation using Phenix revealed a range between 3.0□Å and 5.0□Å across the ArnA complex, with clear secondary structure elements (Sup. Fig. 5B). The model displayed good agreement with the density (Sup. Fig. 5C) and supported by a model-to-map correlation coefficient (CC_mask) of 0.62 using Phenix. The map/model FSC at 0.5 reaches a resolution of 7.5 Å. The overall cryo-EM map resolution, based on the FSC_0.143_ cutoff, was 4.0□Å (Sup. Fig. 5A). The final atomic model comprised 12 chains and 3,726 residues, reflecting the hexameric organization of ArnA. Geometry validation indicated a MolProbity score of 2.36, with a clashscore of 19.09. Ramachandran analysis showed 95.4% of residues in favoured regions and only 0.03% outliers (Table 2).

To assess structural accuracy, the ArnA cryo-EM model was aligned with a previously determined structure (PDB: 6PIH; Yang et al., 2019) using ChimeraX (Meng et al., 2023) (Figure 2C). The overall structural alignment yielded a root-mean-square deviation (RMSD) of 1.2□Å across 183 pruned atom pairs (2.7□Å across all 330 pairs), indicating a high degree of structural similarity.

In terms of domain organization, ArnA is a bifunctional cytoplasmic enzyme composed of two catalytic domains: an N-terminal transformylase domain, which catalyzes the formylation of UDP-4-amino-4-deoxy-L-arabinose (UDP-Ara4N), and a C-terminal dehydrogenase domain, which is responsible for the oxidative conversion of UDP-glucuronic acid., which catalyzes the nicotinamide adenine dinucleotide, oxidized form (NAD^+^)-dependent decarboxylation of UDP-glucuronic acid (Fischer et al., 2015; Gatzeva-Topalova et al., 2005; Yang et al., 2019). These domains are connected by a flexible linker and act sequentially in the Ara4N biosynthesis pathway (Fischer et al., 2015).

Despite the preservation of the global fold, structural comparison revealed several conformational differences relative to the reference model (PDB ID: 6PIH). Notably, loop regions within the transformylase domain, including residues Glu69–Ala98 in Chain D and Pro65–Ser75 in Chain F, exhibited clear deviations (Figure 2C). Additional variations were observed in other surface-exposed loops, suggesting local flexibility or alternative domain orientations that may be influenced by differences in sample conditions or oligomeric assembly. A previous study has shown that ArnA can adopt multiple oligomeric forms, including hexamers and tetramers, and undergoes conformational changes upon substrate binding (Yang et al., 2019). In particular, regions near residues 500–509 and 605–616 have been reported to shift in response to UDP-glucuronic acid, facilitating NAD⁺ coordination (Gatzeva-Topalova et al., 2005).

These observations indicate that the conformational differences identified in the current model, especially within the transformylase domain, may be functionally relevant and reflect the dynamic behavior of ArnA. In addition, the peptide chain of the ArnA model matches well with the experimental density map (Figure 2D).

Altogether, this analysis provides the highest-resolution cryo-EM reconstruction of ArnA at 4.0□Å currently available in the EM and Protein Data Banks, exceeding the resolution of previously published models.

### 3.6. Structural Refinement and Validation of AcrB at Near-Atomic Resolution

Model building for AcrB was carried out using the crystal structure PDB: 7RR7 as the initial template (Trinh et al., 2023). The model was fitted into the 2.92 Å cryo-EM map through molecular replacement using MolRep within the CCP-EM interface (Figure 3A). Refinement was performed using Phenix Real Space Refinement, followed by optimization in Refmac and final adjustments in Coot to complete the atomic model (Figure 3B).

**Figure 3.**
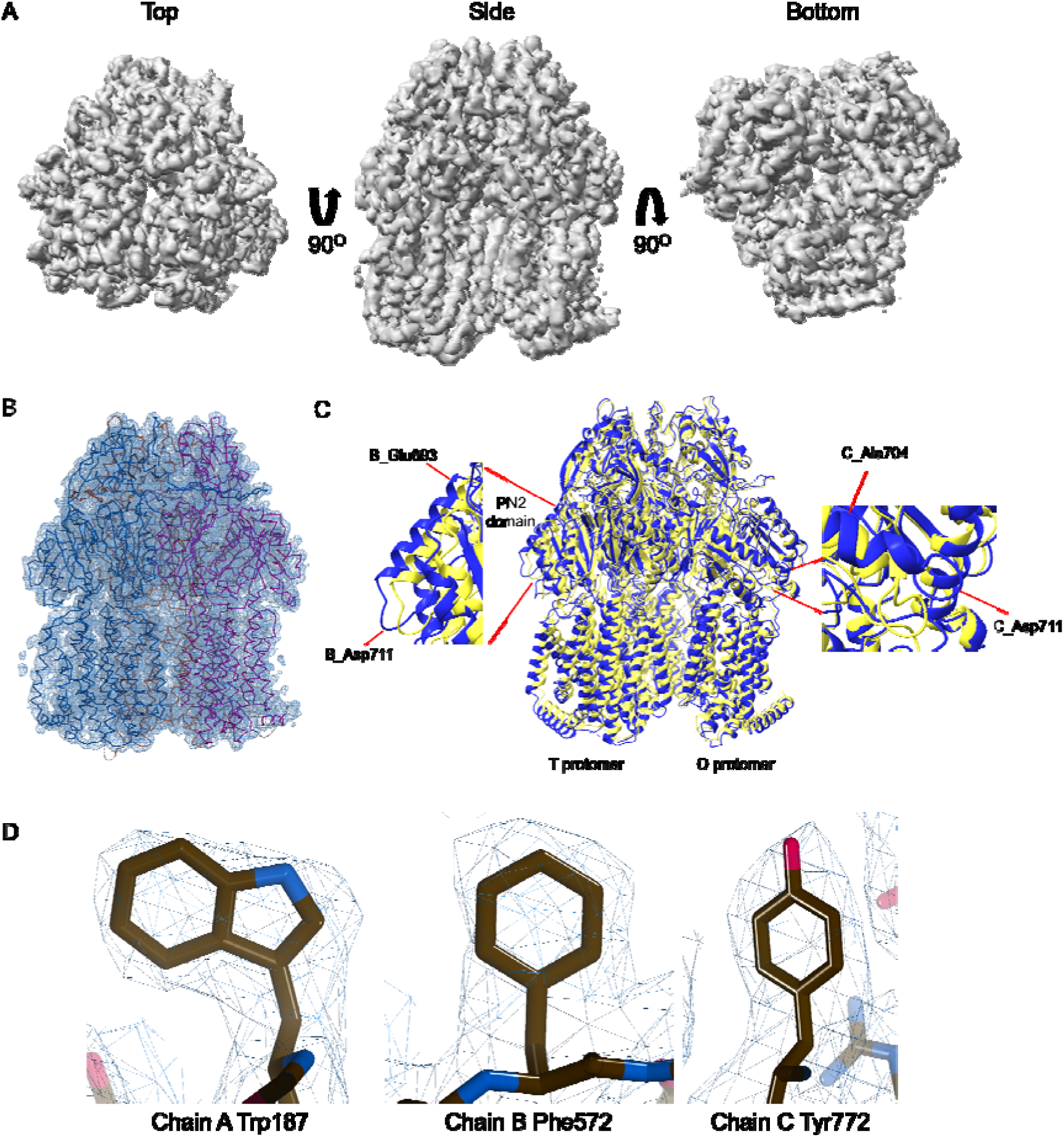
Cryo-EM density map and model fitting of the AcrB trimer. (A) Cryo-EM density map of AcrB reconstructed at 2.92LJÅ resolution, shown from top, side, and bottom views. The map reveals clear secondary structure elements and well-resolved transmembrane helices, consistent with the known trimeric architecture. (B) Final atomic model of AcrB fitted into the cryo-EM density map. Each protomer is rendered in a different color to highlight the asymmetric trimer organization. (C) Structural alignment of the final cryo-EM model (blue) with the reference crystal structure (PDB: 7RR7, yellow), performed using ChimeraX. The RMSD between the two structures is 1.2LJÅ. Zoomed-in views highlight local conformational deviations, including Glu693-Asp711 in protomer B (left) and Ala704-Asp711 in protomer C (right), located within the PN2 subdomain. (D) Close-up view showing the density fit for selected side chains (Trp187 in Chain A, Phe572 in Chain B, Tyr772 in Chain C). The mesh is contoured at 1.6σ in Coot, showing well-defined density for aromatic residues.

The final AcrB model consisted of 3 chains, 3,059 residues, and 23,153 atoms. Model validation metrics indicated high stereochemical quality, with a MolProbity score of 1.95, a clashscore of 10.36, and 97.2% of residues located in favoured Ramachandran regions (Table 2). Notably, no outliers were observed. The model-to-map correlation coefficient (CC_mask) was 0.73 (Table 2). The global cryo-EM map resolution, calculated using the FSC_0.143_ criterion, was 2.92□Å (Sup. Fig. 6A). The map/model FSC at 0.5 reaches a resolution of 3.7 Å (Sup. Fig. 6C). These values collectively reflect a well-refined and accurate model, supported by clearly resolved sidechain features throughout all domains (Figure 4B, 4E).

To evaluate the accuracy of the refined model, the cryo-EM structure of AcrB was compared with the high-resolution crystal structure (PDB: 7RR7); (Trinh et al., 2023). Structural alignment confirmed that the overall trimeric fold was preserved, with an RMSD of 1.2□Å across 759 atom pairs (Figure 3C). Structurally, AcrB functions as a homotrimeric multidrug efflux pump that undergoes a conformational cycling mechanism, with each protomer adopting one of three alternating states: access (loose), binding (tight), or extrusion (open) (Murakami et al., 2002; Seeger et al., 2006). This asymmetric arrangement enables unidirectional substrate transport. In the present model, the trimeric architecture and domain organization are consistent with this established mechanism.

While the global fold aligns well with the reference structure, local differences were identified, particularly in flexible loop regions and surface-exposed side chains. Notable deviations were observed between Asp711 and Glu693, which reside within the PN2 subdomain of the T protomer (Seeger et al., 2006) (Figure 3C). This region has been implicated in substrate accommodation and undergoes conformational shifts during the efflux (Murakami et al., 2002; Ababou & Koronakis, 2016). Furthermore, although the gate loop (residues 615–620) was not directly analyzed in this study, previous structural and mutagenesis studies have shown that this segment modulates the progression of substrates between the proximal and distal binding pockets, especially in the case of large molecules such as erythromycin (Ababou & Koronakis, 2016).

The refined model also shows reasonable side-chain-level agreement with the cryo-EM density in well-ordered regions (Figure 3D), further supporting its accuracy. The data confirm that AcrB at 2.92 Å (Sup. Fig. 5A), although not the intended target of structural analysis, was abundantly and stably present in the FtsH–YidC sample, enabling near-atomic characterization via single-particle cryo-EM (Figure 1).

## 4. DISCUSSION

### 4.1. Aggregation Behaviour Observed in Cryo-EM Compared to NS-EM

Although the same protein samples were used, a clear difference in particle distribution was observed between cryo-EM and negative-stain EM (NS-EM). NS-EM micrographs showed well-dispersed particles and no aggregation. In contrast, aggregation was present in cryo-EM grids, depending on the sample preparation conditions. In the first cryo-EM screening, DDM was used as the main detergent, and the sample was crosslinked with 0.25□mM DSP. CHAPS was not included at this stage, and aggregated particles were evident (not shown).

In the second and third screenings, 0.4% w/v CHAPS was added to the 1 mM DSP-crosslinked sample 10 minutes before vitrification. Under these conditions, particle distribution improved, and no aggregation was observed. CHAPS, a zwitterionic detergent, has been reported to reduce surface tension and support membrane protein stabilization during freezing steps (Li, 2022; Kampjut et al., 2021; Egri et al., 2023). Its use just before vitrification may have helped to maintain sample quality on the grid without affecting detergent micelle formation during purification.

In a separate experiment, CHS was added together with DDM during solubilization and purification. CHS mimics native cholesterol and has been shown to enhance membrane protein integrity in structural studies (Li, 2022; Miyata et al., 2025). Among all conditions tested, the CHS-containing sample displayed the lowest level of aggregation, and grid quality was notably higher (Sup. Fig. 3).

### 4.2. Absence of FtsH–YidC Complex Reconstruction in Cryo-EM Analysis

The aim of our work was to determine the structure of the FtsH–YidC complex using single-particle cryo-EM. However, neither 2D class averages nor 3D classes resembling FtsH, YidC or FtsH–HflKC assemblies were observed in the datasets (Ghanbarpour et al., 2025; Kedrov et al., 2016; Kumazaki et al., 2014; Langklotz et al., 2012, 2012; Lee et al., 2011; Ma et al., 2022; Qiao et al., 2022; Tanaka et al., 2018). Among the expected components, only 2D class averages resembling the cytoplasmic AAA+ domain of FtsH were detected (Figure 1). Side views corresponding to full-length FtsH, i.e. including the transmembrane region, were not observed (Carvalho et al., 2021; Ghanbarpour et al., 2025; W. Liu et al., 2022; Ma et al., 2022; Qiao et al., 2022). FtsH’s transmembrane and cytoplasmic regions are known to adopt variable conformations (Carvalho et al., 2021; W. Liu et al., 2022), and this dynamics likely interfered with consistent alignment and classification during image processing, yielding no side views of FtsH. Additionally, our recent biochemical evidence indicates that the interaction between FtsH and YidC is transient rather than stable, further reducing the likelihood of capturing a complete and well-defined FtsH-YidC complex in vitrified samples (Caliseki et al., 2025)

No particle classes corresponding to full-length YidC were identified either. This may be due to its relatively small size (∼61□kDa), and its flexibility between the transmembrane and periplasmic domains which limit its visibility and alignment in single-particle cryo-EM. Despite strong biochemical evidence for the FtsH-YidC interaction (Sup Fig. 1-3), we could not visualise the complex in single-particle cryo-EM.

### 4.3. High-Resolution Reconstruction of Off-Target Proteins ArnA and AcrB

While the targeted FtsH–YidC complex could not be resolved in cryo-EM datasets, cryo-EM models of two abundant off-target proteins, ArnA and AcrB, were successfully reconstructed at high resolution. ArnA, a cytoplasmic hexamer involved in lipid A modification, was reconstructed at 4.0□Å resolution, and AcrB, a trimeric inner membrane transporter of the AcrAB-TolC efflux system, was resolved at 2.92□Å. Both proteins are known to form stable oligomeric assemblies and are frequently reported as co-purifying species in protein preparations (Andersen et al., 2013; Glover et al., 2011; Veesler et al., 2008).

Mass spectrometry analysis confirmed the presence of ArnA and AcrB in samples purified using both Ni-NTA and Strep-Tactin affinity chromatography (Sup. Table 1) (Caliseki et al., 2025). However, these proteins were not clearly detected in SDS-PAGE analyses of SEC fractions, indicating that their concentrations at this stage were below the sensitivity of gel-based visualization. This difference between MS and SDS-PAGE results suggests that while their overall abundance was low, their particle abundance in cryo-grids, structural rigidity and symmetry were sufficient to enable efficient classification and reconstruction. In addition, their particle diameter is similar to cytoplasmic domain of FtsH (14 Å).

ArnA is a cytoplasmic protein and would not be expected to co-enrich with membrane fractions in ultracentrifugation. Its identification in detergent-solubilized membrane protein samples may indicate a possible interaction, directly or indirectly, with membrane components such as FtsH or YidC. Despite being below the detection threshold in Coomassie-stained SDS gels, ArnA yielded a 4.0□Å cryo-EM map, which currently represents the highest-resolution cryo-EM structure of this protein available in the Electron Microscopy Data Bank (EMDB), and also in the PDB.

Similarly, AcrB was consistently identified in mass spectrometry across different purification steps (Sup. Table 1) (Caliseki et al., 2025), reflecting its structural integrity and biochemical stability in detergent micelles. These properties likely facilitated its retention through the cryo-EM grid preparation and image processing workflow facilitated by accurate particle alignment (Su et al., 2021).

### 4.4. Cryo-EM Reveals Co-Purified Components Linked to Proteostasis and Aerobic Metabolism

In this study, FtsH, YidC, and HflKC were co-overexpressed; however, purification predominantly yielded FtsH and YidC, while HflKC was not reliably detected. Two factors may explain the absence of HflKC. First, the mutually exclusive interaction between HflKC and YidC for FtsH binding could lead to competitive exclusion of HflKC. Second, previous transcriptomic studies have shown that depletion of YidC leads to increased expression of HflK and HflC (Wickström et al., 2011), raising the possibility that YidC overexpression may inversely affect HflKC levels. However, further experiments would be required to directly assess HflKC expression levels or membrane incorporation under these conditions. Cryo-EM 2D class averages revealed particles corresponding to cytochrome bo₃ oxidase, a key component of the aerobic respiratory chain that requires YidC for membrane insertion (Celebi et al., 2006; Du Plessis et al., 2006). This observation may reflect a functional association with YidC, consistent with its known role in respiratory complex biogenesis. Alternatively, the presence of cytochrome bo₃ may be influenced by the apparent absence or very low abundance of HflKC, which remained barely detectable in the purified sample despite being co-overexpressed (Caliseki et al., 2025). HflKC has been implicated in the regulation of cytochrome oxidase expression and aerobic metabolism (Perez-Lopez et al., 2025). Although this remains speculative in the context of the present study, the findings raise the possibility that HflKC levels may influence the abundance or stability of cytochrome bo₃ under stress conditions.

Finally, the failure to observe an intact FtsH–YidC complex, along with the enrichment of cytochrome bo₃ (Borisov et al., 2021) and GroEL (Fourie & Wilson, 2020), likely reflects the dynamic and transient nature of YidC interactions, and the interactome being shaped by the cellular metabolic state and proteostasis demands. The complexity of these interactions is further reflected in the difficulty of resolving intact YidC complexes in our cryo-EM datasets. However, recent advances, such as the BaR method, have demonstrated the capability of Cryo-EM to resolve multiple protein structures from heterogeneous samples (Su et al., 2021). The BaR approach highlights Cryo-EM’s remarkable power to capture diverse structural states within a single dataset, even from complex mixtures. In our study, this capability allowed us to resolve the structures of ArnA and AcrB while also identifying 2D class averages resembling GroEL and cytochrome bo₃, further emphasizing the dynamic nature of the protein mixtures.

In summary, our findings underscore the complexity of cryo-EM datasets derived from membrane protein complexes and highlight the importance of integrating structural and biochemical data with mass spectrometry to accurately interpret the data.

## 5. CONCLUSION

This study highlights the complexity of transient membrane protein interactions, such as YidC and FtsH involved in membrane protein homeostasis, and the associated challenges to obtain structural and functional insights into these protein complexes by cryo-EM. While the FtsH–YidC complex could not be reconstructed, high-resolution structures of off-target proteins, such as ArnA and AcrB, were successfully obtained, providing valuable structural insights into these components. The enrichment of co-purified proteins in cryo-EM 2D class averages, such as GroEL and cytochrome bo₃ (identified also in mass spectrometry) further illustrates the complexity of the task. Our findings emphasize the importance of combining structural and biochemical approaches and mass spectrometry to gain a comprehensive understanding of membrane protein function and regulation, particularly in the context of stress-induced cellular responses.

## Acknowledgements

M.Ç., and B.V.K. were funded by the Scientific and Technological Council of Türkiye (TÜBİTAK) BİDEB 2232 International Outstanding Researchers Program (Project No: 118C225). M.Ç. was also funded by TÜBİTAK 2211-A National Graduate Fellowship Program. C.S. acknowledges funding by a BBSRC Responsive Mode Grant (BB/P000940/1) and a Wellcome Trust Investigator Grant (210701/Z/18/Z). M.Ç. was also supported by a TÜBİTAK 2214-A - International Research Fellowship Program for carrying out a part of his PhD thesis studies at the University of Bristol, School of Biochemistry. We acknowledge Dr. Süreyya Özcan and Hatice Akkulak from METU for their help in XL-MS analyses. We acknowledge support and assistance by the Wolfson Bioimaging Facility and the GW4 Facility for High-Resolution Electron Cryo-Microscopy funded by the Wellcome Trust (202904/Z/16/Z and 206181/Z/17/Z) and BBSRC (BB/R000484/1).

## Contributions

M.Ç., C.S., and B.V.K. conceived the study. M.Ç. performed all experiments and wrote the manuscript with input from C.S. and B.V.K. U.B. contributed to Cryo-EM sample vitrification and data collection. S.K.N.Y. contributed to NS-EM analysis and initial Cryo-EM data processing. All authors have read and agreed to the final manuscript.

## Data Availability Statement

All data needed to evaluate the conclusions in the paper are present in the paper and/or the Supplementary Materials. All datasets generated during the current study have been deposited in the Electron Microscopy Data Bank (EMDB) under accession numbers EMD-64789 (ArnA), EMD-64793 (AcrB), and in the Protein Data Bank (PDB) under accession number PBD ID 9V5H (ArnA) and 9V5R (AcrB).

---

**Sup. Fig 1.**
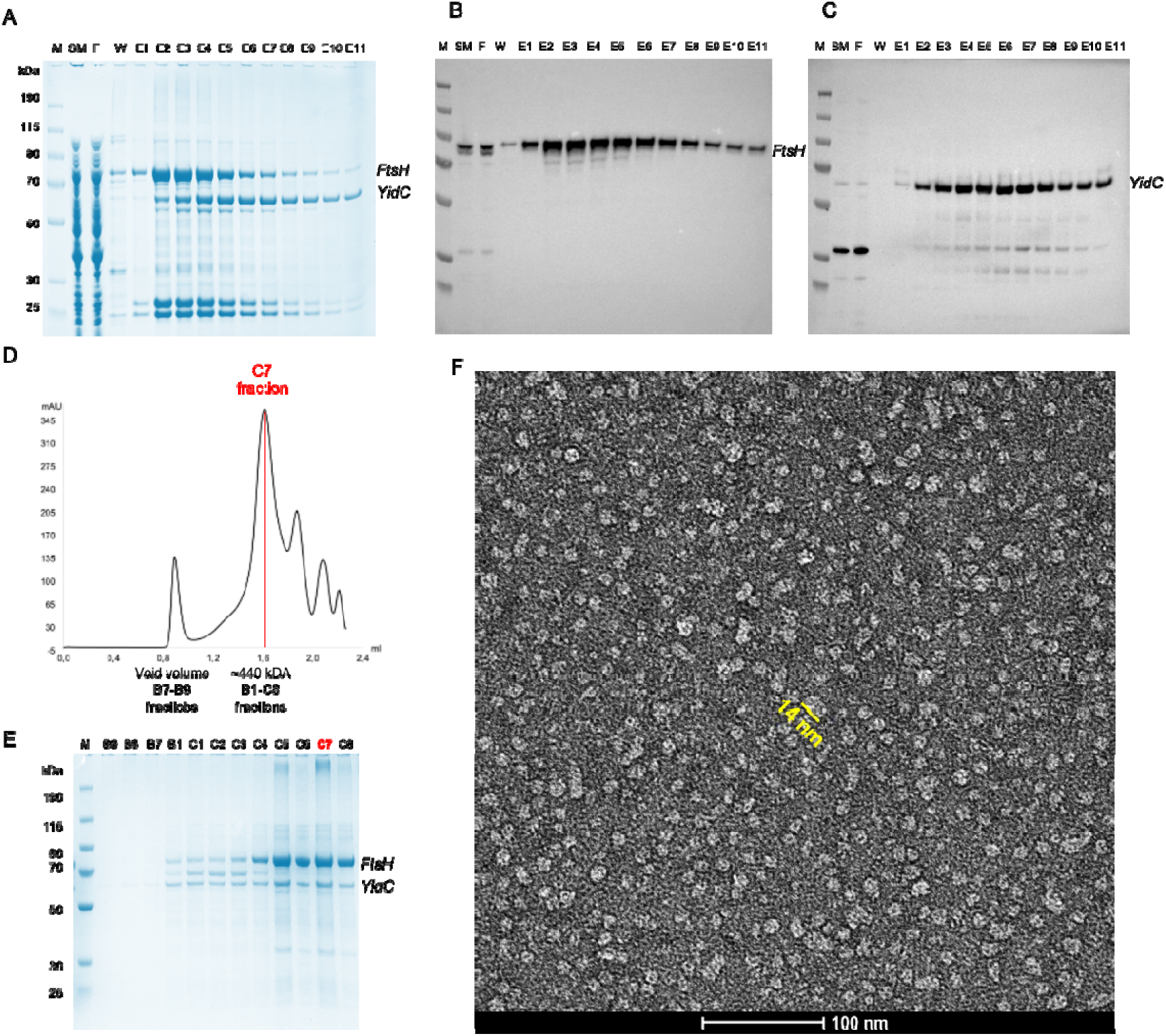
Purification of the FtsH–YidC sample with only DDM. (A) Coomassie stained SDS gel of Ni-NTA purified samples. (B) Western blot using anti-StrepTag II antibody to detect the presence of FtsH. (C) Western blot using anti-His antibody to detect His-tagged YidC. (D) Superose 6 Increase 3.2/300 SEC chromatogram from ÄKTAmicro. Fraction C7 was further analysed. (E) Coomassie-stained SDS gel of SEC fractions corresponding to first and second peak. (F) Negative-stain EM micrograph of C7 fraction. Scale bar of a particle: 14 nm.

**Sup. Fig 2.**
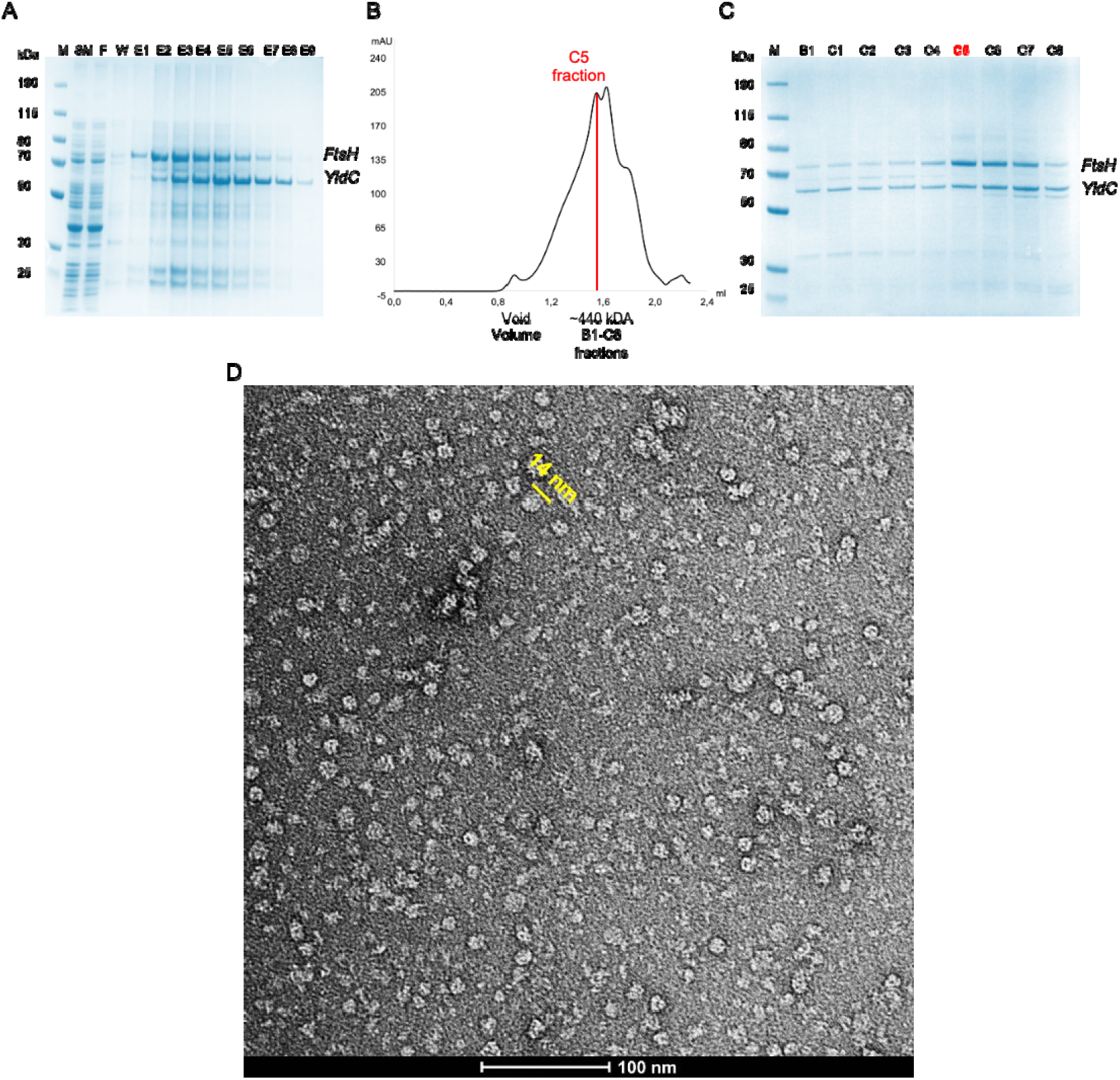
Purification of crosslinked FtsH–YidC sample with DDM-CHAPS combination. (A) Coomassie-stained SDS gel of Ni-NTA purified samples. (B) Superose 6 Increase 3.2/300 SEC chromatogram from ÄKTAmicro. Fraction C5 was further analysed. (C) Coomassie-stained SDS gel of SEC fractions. (D) Negative-stain EM micrograph of C5 fraction. Scale bar of a particle: 14 nm.

**Sup. Fig 3.**
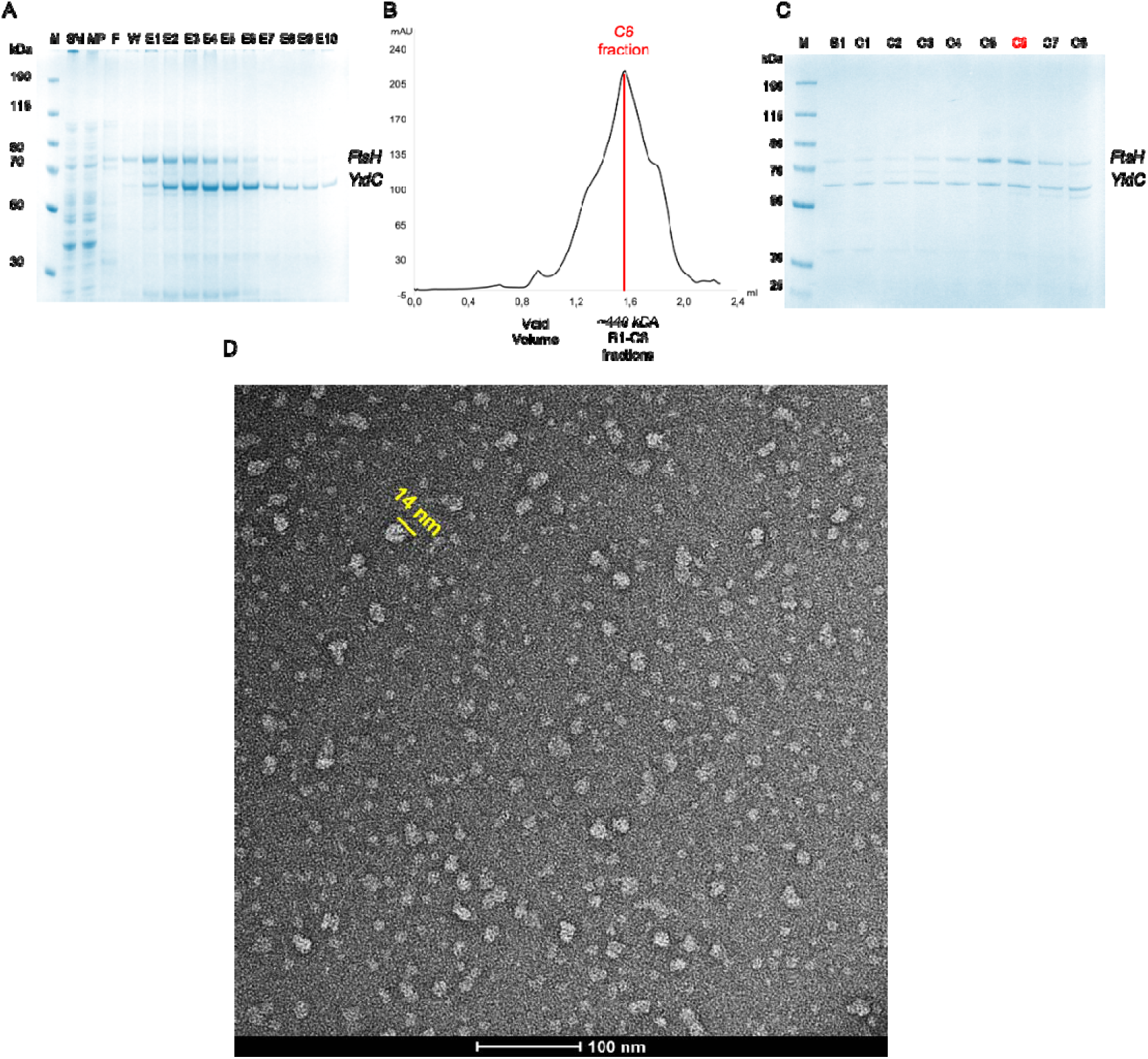
Purification of crosslinked FtsH–YidC sample with DDM-CHS combination. (A) Coomassie-stained SDS gel of Ni-NTA purified samples. (B) Superose 6 Increase 3.2/300 SEC chromatogram. Fraction C6 was further analysed. (C) Coomassie-stained SDS gel of SEC fractions. (D) Negative-stain EM micrograph of C5 fraction. Scale bar of a particle: 14 nm.

**Supplementary Figure 4.**
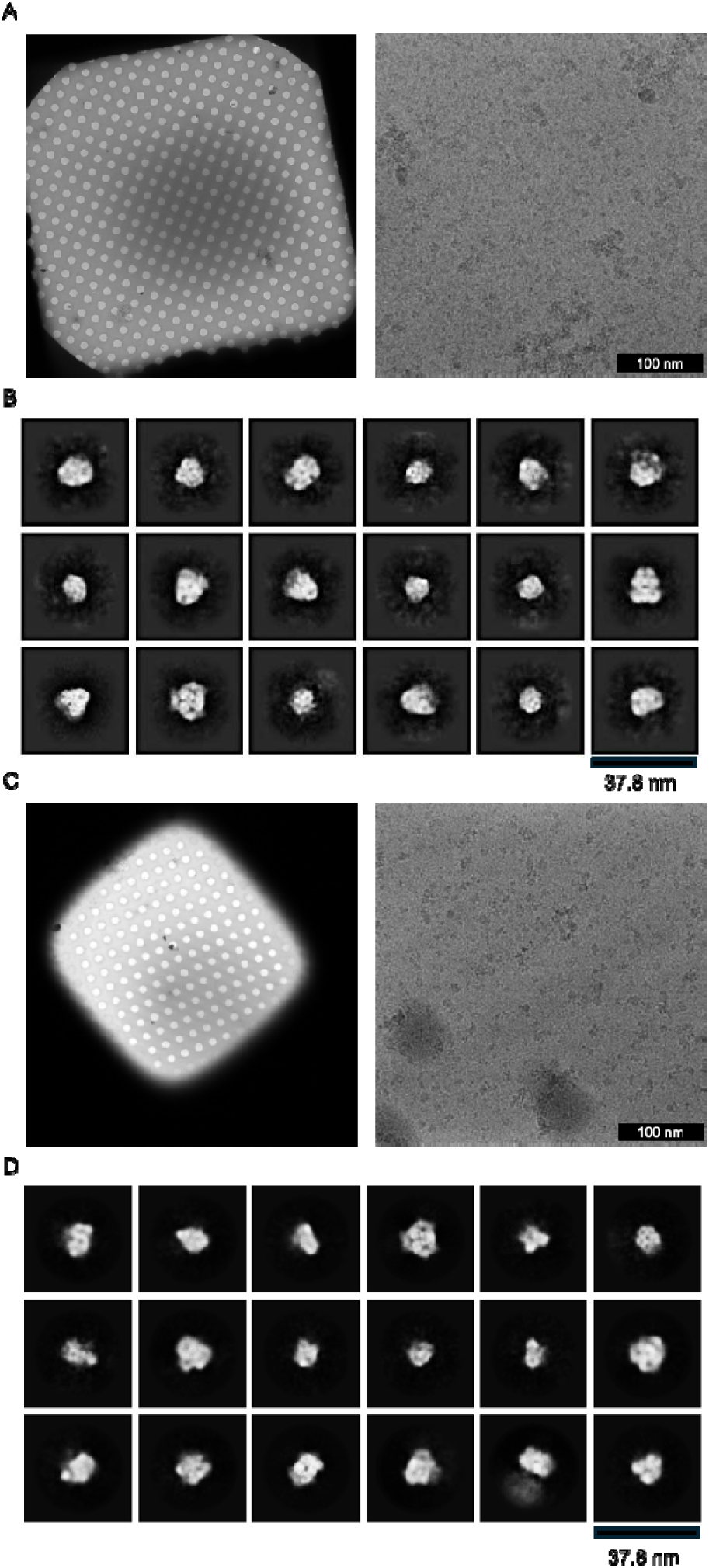
Cryo-EM screening of purified FtsH–YidC samples. (A) Micrograph of sample using C5 fraction after SEC with 0.4% w/v CHAPS as an additive prior to vitrification. Screening revealed dispersed particles suitable for further image analysis. (B) Optimized micrograph after applying DDM-CHS combination for solubilization and purification. C6 fraction from SEC was used. Screening provided high-quality micrographs with well-dispersed particles suitable for data acquisition and downstream analysis. Scale bar: 14 nm.

**Supplementary Figure 5.**
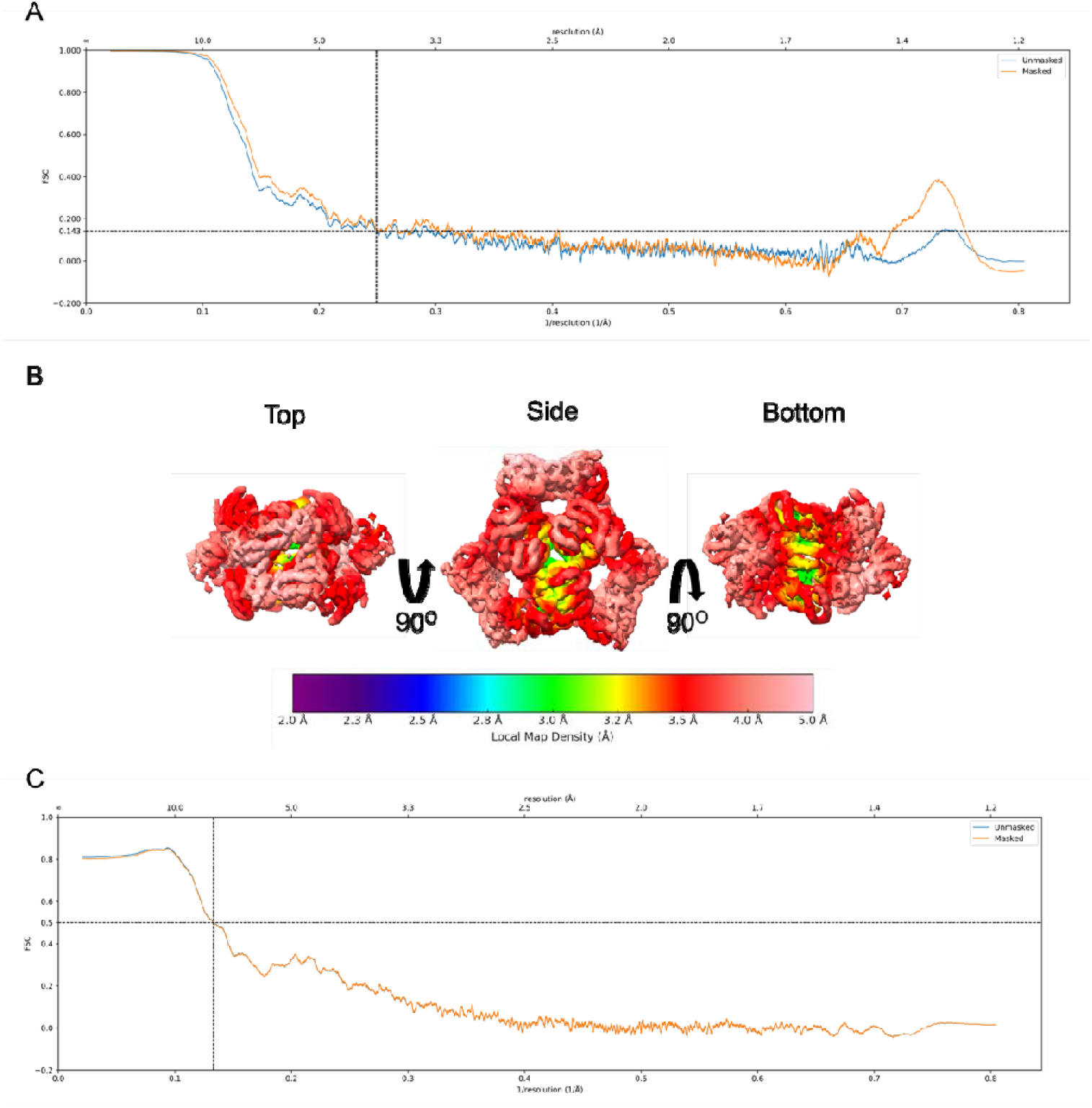
Quality of the ArnA map and model. (A) The Fourier Shell Correlation (FSC) curve for ArnA after refinement of ArnA particles. The FSC 0.143 criterion indicates an overall resolution of 4.0 Å. Blue curve: FSC curve of unmasked maps; orange curve: FSC curve of phase randomized masked maps. (B) Local resolution of the final ArnA cryo-EM map calculated in PHENIX. The core of the complex is resolved at 3.5 Å whereas peripheral parts have a lower resolution of ∼ 4 Å. (C) FSC curve calculated between the ArnA atomic model and the final cryo-EM map. The map/model FSC at 0.5 reaches a resolution of 7.5 Å.

**Supplementary Figure 6.**
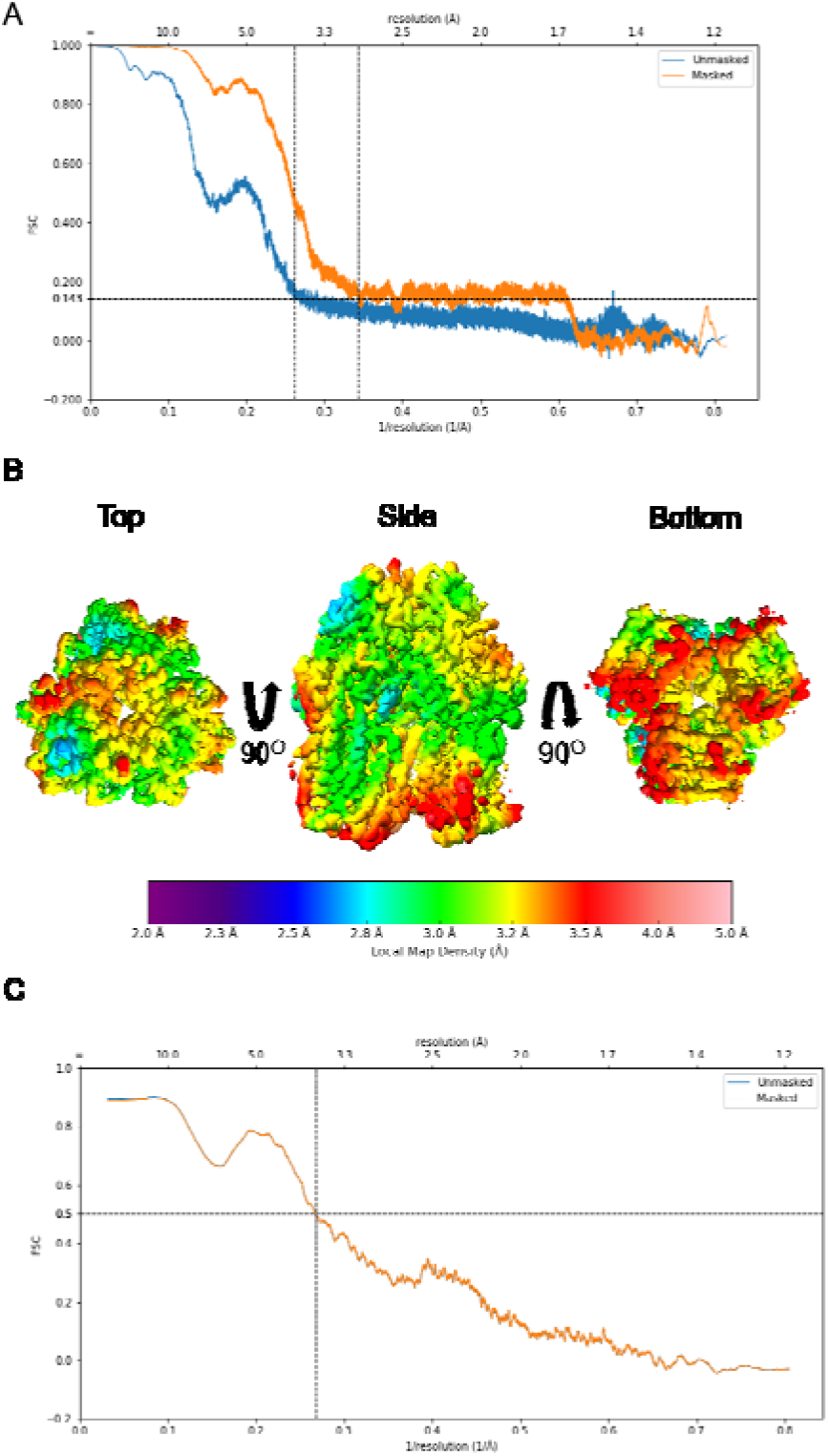
Quality of the AcrB map and model. (A) The Fourier Shell Correlation (FSC) curve for AcrB after gold-standard refinement of AcrB particles (blue curve). The FSC = 0.143 criterion indicates an overall resolution of 2.92 Å. Blue curve: FSC curve of unmasked maps; orange curve: FSC curve of phase randomized masked maps. (B) Local resolution of the final ArnA cryo-EM map calculated in PHENIX. The core of the complex is resolved at 3 Å whereas peripheral parts have a lower resolution of ∼ 3.5 Å. (C) FSC curve calculated between the ArnA atomic model and the final cryo-EM map. The map/model FSC at 0.5 reaches a resolution of 3.7 Å.

**Supplementary Table 1.**
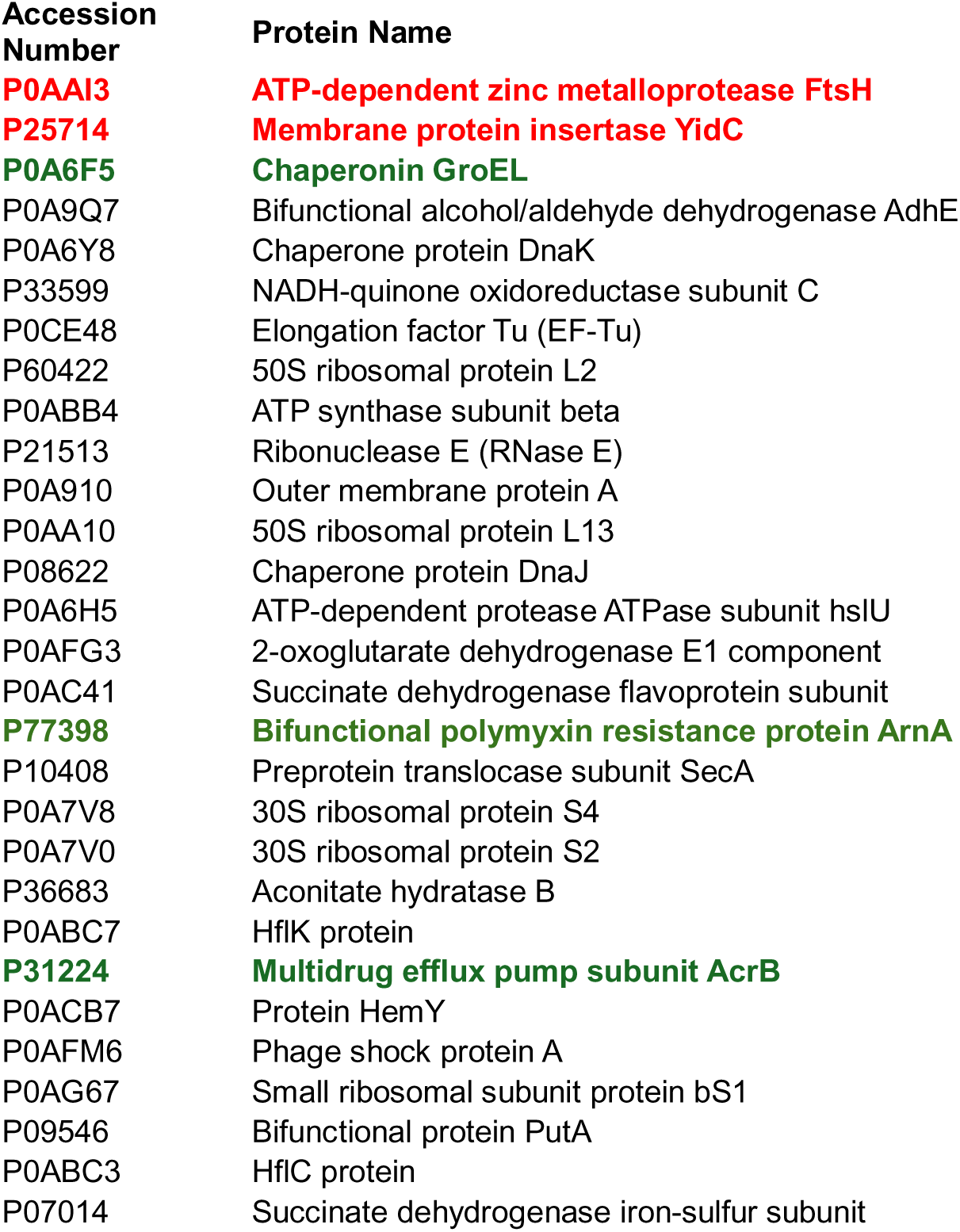
Proteins co-purified with FtsH and YidC. MS analysis of DSP-treated, SEC-purified FtsH-YidC sample purified from an *E. coli* C43(DE3) strain overexpressing FtsH, YidC, HflK and HflC (Caliseki et al., 2025). Target proteins are in red colour, off-target proteins detected in cryo-EM 2D class averages in green colour.

