## Supplementary Information for "Off-Target Structural Insights: ArnA and AcrB in Bacterial Membrane Protein Cryo-EM Analysis"

**Supplemental Information**

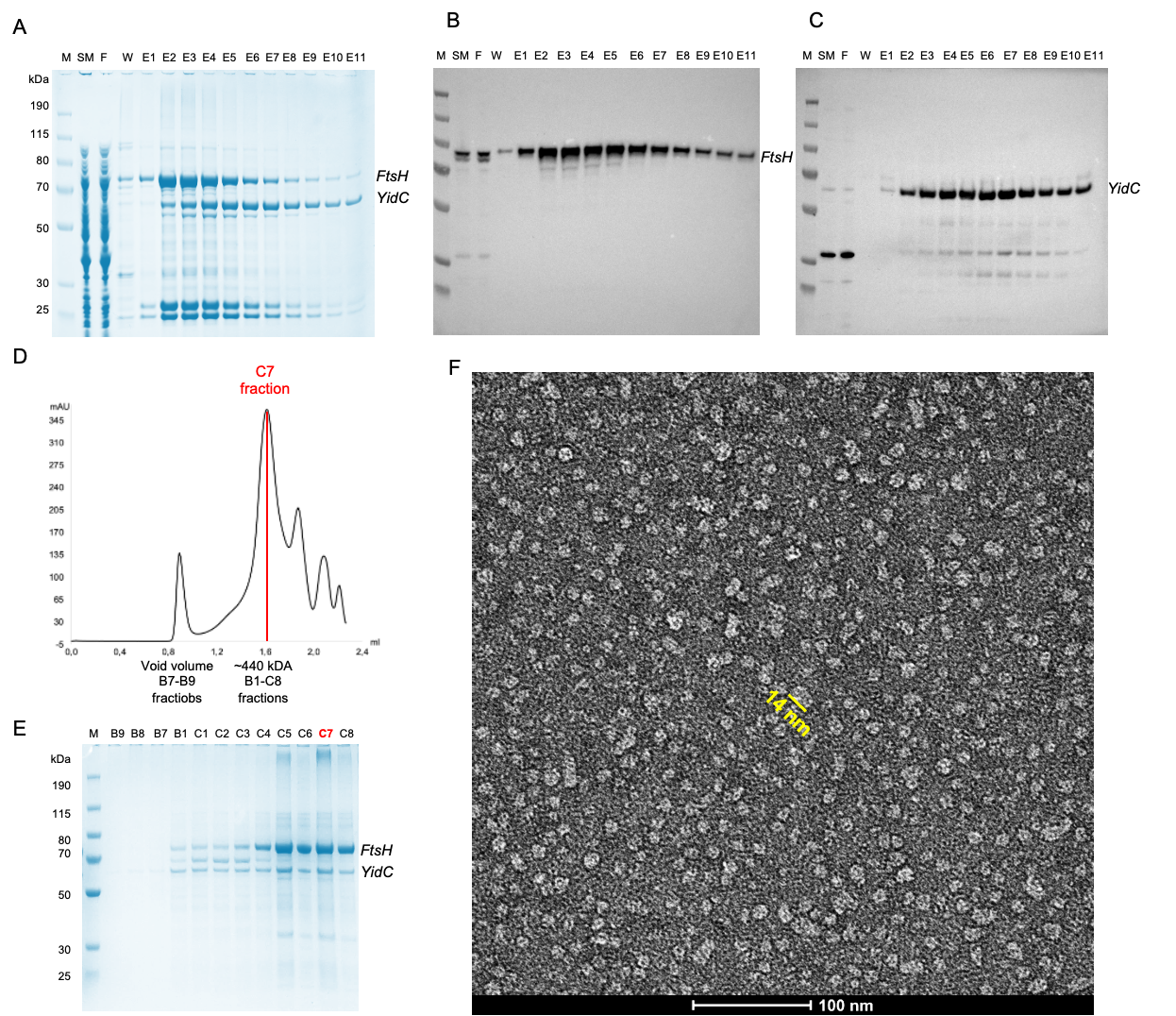

**Sup. Fig 1. Purification of the FtsH–YidC sample with only DDM.** (A) Coomassie stained SDS gel of Ni-NTA purified samples. (B) Western blot using anti-StrepTag II antibody to detect the presence of FtsH. (C) Western blot using anti-His antibody to detect His-tagged YidC. (D) Superose 6 Increase 3.2/300 SEC chromatogram from ÄKTAmicro. Fraction C7 was further analysed. (E) Coomassie-stained SDS gel of SEC fractions corresponding to first and second peak. (F) Negative-stain EM micrograph of C7 fraction. Scale bar of a particle: 14 nm.

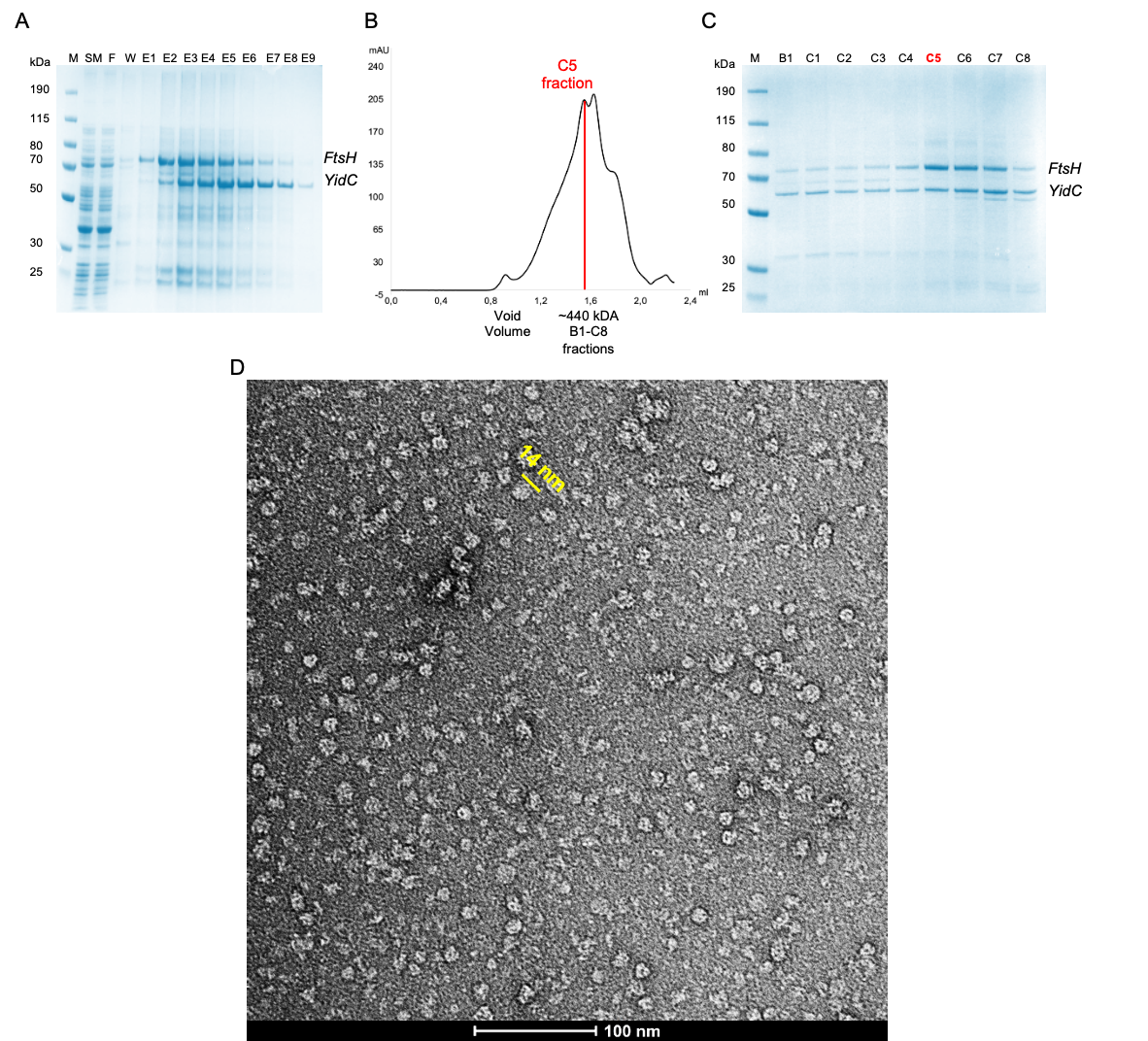

**Sup. Fig 2. Purification of crosslinked FtsH–YidC sample with DDM-CHAPS combination.** (A) Coomassie-stained SDS gel of Ni-NTA purified samples. (B) Superose 6 Increase 3.2/300 SEC chromatogram from ÄKTAmicro. Fraction C5 was further analysed. (C) Coomassie-stained SDS gel of SEC fractions. (D) Negative-stain EM micrograph of C5 fraction. Scale bar of a particle: 14 nm.

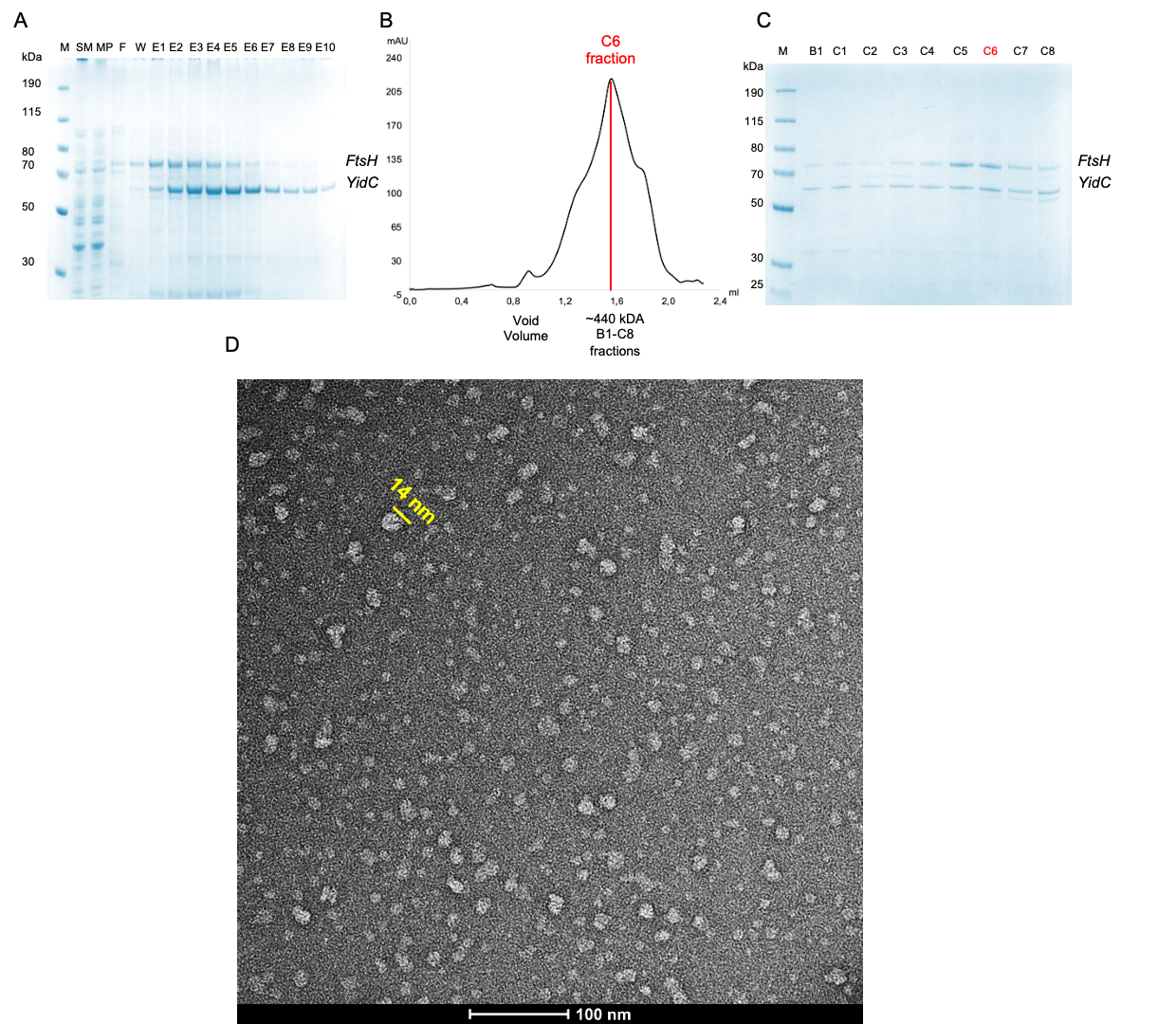

**Sup. Fig 3. Purification of crosslinked FtsH–YidC sample with DDM-CHS combination.** (A) Coomassie-stained SDS gel of Ni-NTA purified samples. (B) Superose 6 Increase 3.2/300 SEC chromatogram. Fraction C6 was further analysed. (C) Coomassie-stained SDS gel of SEC fractions. (D) Negative-stain EM micrograph of C5 fraction. Scale bar of a particle: 14 nm.

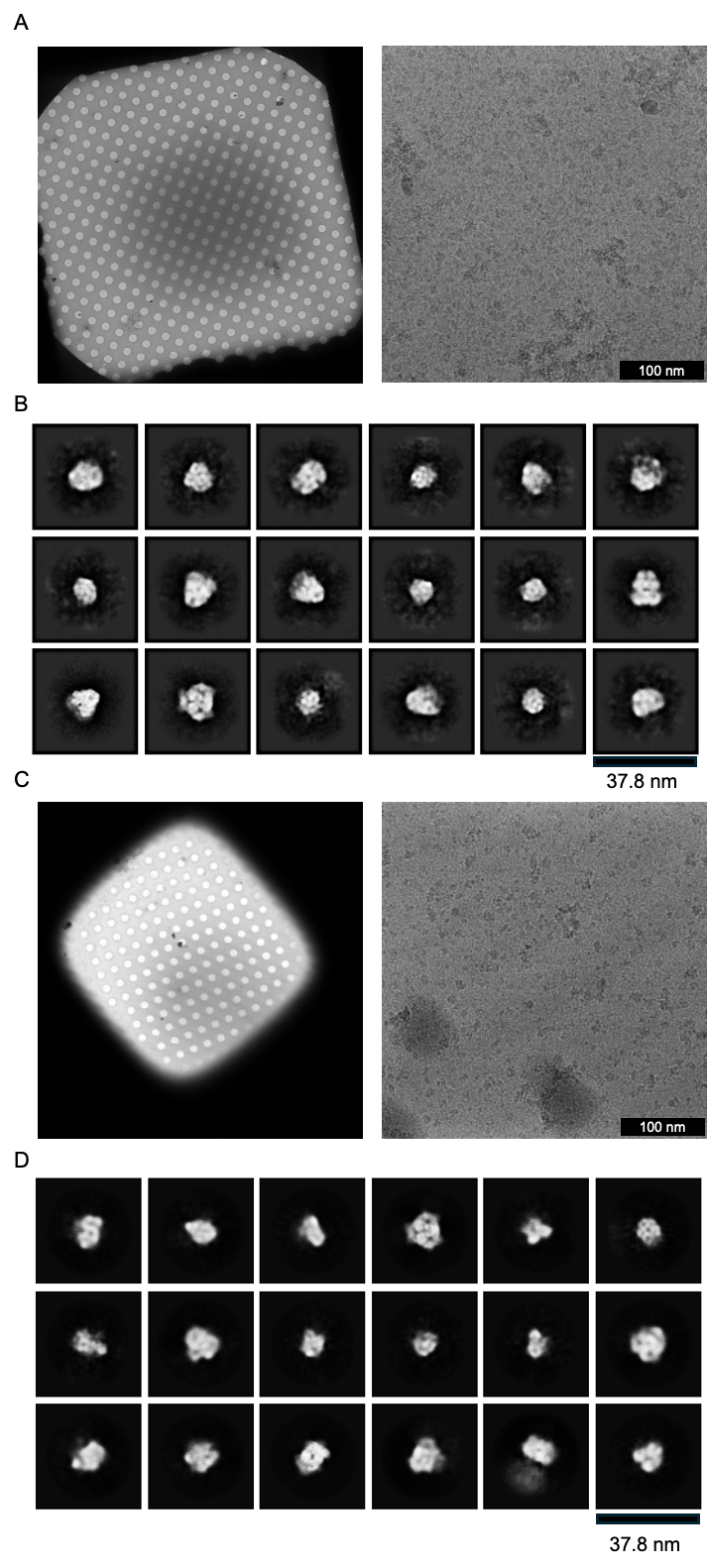

**Supplementary Figure 4. Cryo-EM screening of purified FtsH–YidC samples.** (A) Micrograph of sample using C5 fraction after SEC with 0.4% w/v CHAPS as an additive prior to vitrification. Screening revealed dispersed particles suitable for further image analysis. (B) Optimized micrograph after applying DDM-CHS combination for solubilization and purification. C6 fraction from SEC was used. Screening provided high-quality micrographs with well-dispersed particles suitable for data acquisition and downstream analysis. Scale bar: 14 nm.

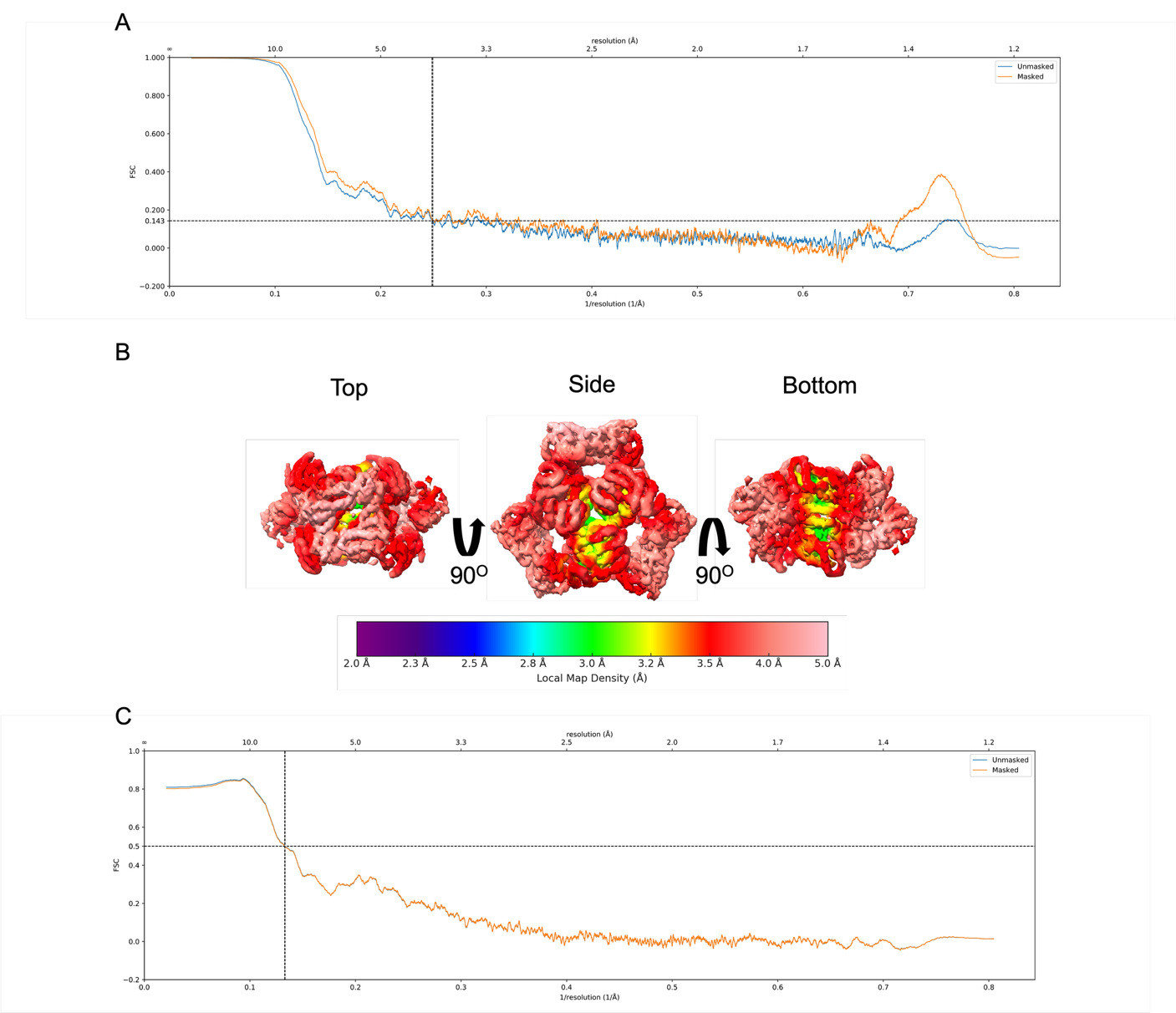

**Supplementary Figure 5.** Quality of the ArnA map and model. (A) The Fourier Shell Correlation (FSC) curve for ArnA after refinement of ArnA particles. The FSC 0.143 criterion indicates an overall resolution of 4.0 Å. Blue curve: FSC curve of unmasked maps; orange curve: FSC curve of phase randomized masked maps. (B) Local resolution of the final ArnA cryo-EM map calculated in PHENIX. The core of the complex is resolved at 3.5 Å whereas peripheral parts have a lower resolution of ~ 4 Å. (C) FSC curve calculated between the ArnA atomic model and the final cryo-EM map. The map/model FSC at 0.5 reaches a resolution of 7.5 Å.

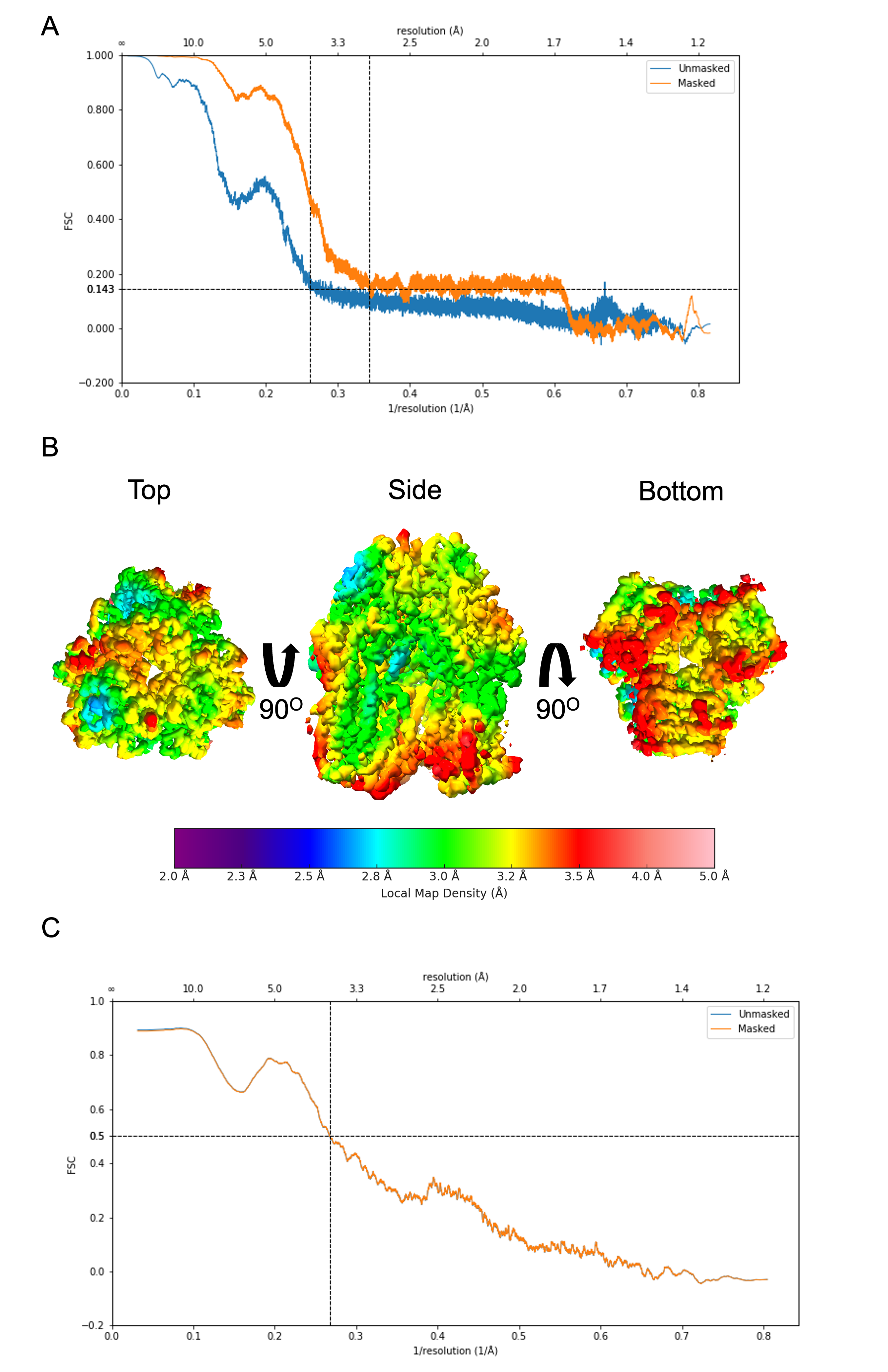

**Supplementary Figure 6.** Quality of the AcrB map and model. (A) The Fourier Shell Correlation (FSC) curve for AcrB after gold-standard refinement of AcrB particles (blue curve). The FSC = 0.143 criterion indicates an overall resolution of 2.92 Å. Blue curve: FSC curve of unmasked maps; orange curve: FSC curve of phase randomized masked maps. (B) Local resolution of the final ArnA cryo-EM map calculated in PHENIX. The core of the complex is resolved at 3 Å whereas peripheral parts have a lower resolution of ~ 3.5 Å. (C) FSC curve calculated between the ArnA atomic model and the final cryo-EM map. The map/model FSC at 0.5 reaches a resolution of 3.7 Å.

**Supplementary Table 1. Proteins co-purified with FtsH and YidC.** MS analysis of DSP-treated, SEC-purified FtsH-YidC sample purified from an *E. coli* C43(DE3) strain overexpressing FtsH, YidC, HflK and HflC (Caliseki et al., 2025). Target proteins are in red colour, off-target proteins detected in cryo-EM 2D class averages in green colour.

| **Accession Number** | **Protein Name** |
| --- | --- |
| **P0AAI3** | **ATP-dependent zinc metalloprotease FtsH** |
| **P25714** | **Membrane protein insertase YidC** |
| **P0A6F5** | **Chaperonin GroEL** |
| P0A9Q7 | Bifunctional alcohol/aldehyde dehydrogenase AdhE |
| P0A6Y8 | Chaperone protein DnaK |
| P33599 | NADH-quinone oxidoreductase subunit C |
| P0CE48 | Elongation factor Tu (EF-Tu) |
| P60422 | 50S ribosomal protein L2 |
| P0ABB4 | ATP synthase subunit beta |
| P21513 | Ribonuclease E (RNase E) |
| P0A910 | Outer membrane protein A |
| P0AA10 | 50S ribosomal protein L13 |
| P08622 | Chaperone protein DnaJ |
| P0A6H5 | ATP-dependent protease ATPase subunit hslU |
| P0AFG3 | 2-oxoglutarate dehydrogenase E1 component |
| P0AC41 | Succinate dehydrogenase flavoprotein subunit |
| **P77398** | **Bifunctional polymyxin resistance protein ArnA** |
| P10408 | Preprotein translocase subunit SecA |
| P0A7V8 | 30S ribosomal protein S4 |
| P0A7V0 | 30S ribosomal protein S2 |
| P36683 | Aconitate hydratase B |
| P0ABC7 | HflK protein |
| **P31224** | **Multidrug efflux pump subunit AcrB** |
| P0ACB7 | Protein HemY |
| P0AFM6 | Phage shock protein A |
| P0AG67 | Small ribosomal subunit protein bS1 |
| P09546 | Bifunctional protein PutA |
| P0ABC3 | HflC protein |
| P07014 | Succinate dehydrogenase iron-sulfur subunit |
